# High sensitivity limited material proteomics empowered by data-independent acquisition on linear ion traps

**DOI:** 10.1101/2022.06.27.497681

**Authors:** Teeradon Phlairaharn, Samuel Grégoire, Lukas R. Woltereck, Valdemaras Petrosius, Benjamin Furtwängler, Brian C. Searle, Erwin M. Schoof

## Abstract

In recent years, the concept of cell heterogeneity in biology has gained increasing attention, concomitant with a push towards technologies capable of resolving such biological complexity at the molecular level. For single-cell proteomics using Mass Spectrometry (scMS) and low-input proteomics experiments, the sensitivity of an orbitrap mass analyzer can sometimes be limiting. Therefore, low-input proteomics and scMS could benefit from linear ion traps, which provide faster scanning speeds and higher sensitivity than an orbitrap mass analyzer, however at the cost of resolution. We optimized an acquisition method that combines the orbitrap and linear ion trap, as implemented on a tribrid instrument, while taking advantage of the high-field asymmetric waveform ion mobility spectrometry (FAIMS) pro interface, with a prime focus on low-input applications. First, we compared the performance of orbitrap-versus linear ion trap mass analyzers. Subsequently, we optimized critical method parameters for low-input measurement by data-independent acquisition (DIA) on the linear ion trap mass analyzer. We conclude that linear ion traps mass analyzers combined with FAIMS and Whisper™ flow chromatography are well-tailored for low-input proteomics experiments, and can simultaneously increase the throughput and sensitivity of large-scale proteomics experiments where limited material is available, such as clinical samples and cellular sub-populations.

## INTRODUCTION

Deep proteome profiling of single cells and low-input material is an attractive discovery-based, hypothesis-free pipeline to study biological heterogeneity in health and diseases. Shotgun proteomics has achieved initial milestones using the latest advances in nanoflow liquid chromatography-tandem mass spectrometry (LC-MS/MS)^1–5^. However, technical challenges remain in the study of scMS and low-input approaches. A variety of fields like sample preparation, experimental throughput, instrument sensitivity, and computational tools still require further optimization^6^.

To maximize the efficiency of low-input proteomics experiments, many sample processing methods were developed to minimize sample losses and surface contact during sample isolation, preparation, and transfer^7–11^. While deep proteome profiling comes with the tradeoff that it requires a longer LC gradient, we here use a Whisper™ stepped pre-formed beta gradient^2^ eluting the peptides at a 100 nL/min flow rate and a 40 or 20 “samples-per-day” (SPD) method to balance robustness and sensitivity. Brunner *et al*.^5^ included a similar platform during their recent demonstration of advances in instrument development to analyze single-cell proteomes on a trapped ion mobility mass spectrometer with diaPASEF^12^. Currently, most single-cell proteomics and low-input proteomics studies were performed on an orbitrap mass analyser instrument^4,13–17^ or a time-of-flight instrument^5,15,18^. Linear ion traps stand as an attractive alternative mass analyzer to orbitraps for low-input applications in mass spectrometry-based proteomics^19^ thanks to their increased sensitivity and efficient scanning speed. To enhance the results of data acquisition for low-input quantitative proteomics experiments, data-independent acquisition (DIA) is an attractive approach, as precursor ions are fragmented and acquired independently from their intensity, which makes this acquisition method less biased and reduces missing values compared to data-dependent acquisition (DDA)^20^.

Here we present LIT-DIA, a data acquisition method that combines the utilization of an orbitrap (OT) for high-resolution MS1 scans, with the linear ion trap (LIT) for low-resolution, but high-sensitivity, MS2 scans^21,22^. We also integrate FAIMSPro ion mobility, which has been shown to decrease chemical background noise, increase specificity^23^, and improve protein coverage in low-input proteomics experiments^24^. We demonstrate the utility of using the LIT as a mass analyzer for ultra-low input samples, simulated by carefully controlled dilution series, and determine at which sample load the LIT starts outperforming the OT. We evaluate the impact of gradient length, DIA window schemes, and MS2 ion injection times (IITs), and present an evaluation of these parameters. This work provides a resource for a comprehensive assessment of LIT-DIA, allowing researchers to implement tribrid instruments in their own DIA-based interrogation of low-input biological samples.

## EXPERIMENT PROCEDURES

### HeLa culture and sample preparation

HeLa cells (ECACC: 93021013, Sigma-Aldrich) were maintained in an H_2_O-saturated atmosphere in Gibco Advanced DMEM (Thermo Fisher Scientific) supplemented with 10% FBS (Gibco, Thermo Fisher Scientific), GlutaMax (Gibco, Thermo Fisher Scientific) and penicillin-streptomycin (Gibco, Thermo Fisher Scientific) at 100 μg/ml at 37 □C and 5% CO_2_. Cells were passaged at 90% confluency by removing the medium, washing with DPBS (Gibco, Thermo Fisher Scientific), and detaching the cells with 2.5 ml of Accutase solution (Gibco, Thermo Fisher Scientific).

Cells were harvested at 80% confluence and lysed in 5% sodium dodecyl sulfate (SDS), 50 mM Tris (pH 8), 75 mM NaCl, and protease inhibitors (Roche, Basel, Switzerland, Complete-mini EDTA-free). The cell lysate was sonicated for 2 × 30 s and then was incubated for 10 minutes on ice. Proteins were reduced and alkylated with 5 mM TCEP and 10 mM CAA for 20 minutes at 45 °C. Proteins were diluted to 1% SDS and digested with MS grade trypsin protease and Lys-C protease (Pierce, Thermo Fisher Scientific) overnight at an estimated 1:100 enzyme to substrate ratio quenching with 1% trifluoroacetic acid (TFA) in isopropanol. For the clean-up step^25^ by styrene-divinylbenzene reverse-phase sulfonate (SDB-RPS), 10 μg of peptides were loaded on StageTip^26^ and washed twice by adding 100 μl of 1% TFA in isopropanol. Peptides were eluted by adding 50 μl of elution buffer (1% Ammonia, 19% ddH_2_O, and 80% Acetonitrile) in a PCR tube and dried at 45 °C in a SpeedVac. Lastly, peptides were resuspended in buffer A and their concentration was measured by nanodrop.

C18 EvoTips were activated by adding 25 μL buffer B (80% Acetonitrile, 0.1% Formic acid), centrifuged at 700 xg and bathed in isopropanol for 1 minute. Then, 50 μL buffer A (0.1% Formic acid) was added to each EvoTip followed by centrifugation at 700 xg for 1 min. The sample was loaded into the EvoTip, followed by centrifugation at 700 xg for 1 min, and two centrifugation steps after adding 50 μL buffer A. Lastly, 150 μL buffer A was added to each EvoTip and spun for 10 sec at 300 xg.

### LC-MS/MS analysis

For all proteome analyses, we used an EvoSep One liquid chromatography system and analyzed the benchmark experiments with diluted tryptic HeLa with a 31- and 58 minutes stepped pre-formed gradient eluting the peptides at a 100 nL/min flow rate. We use a 15 cm x 75 μm ID column (PepSep) with 1.9 μm C18 beads (Dr. Maisch, Germany) and a 10 μm ID silica electrospray emitter (PepSep). Mobile phases A and B were 0.1% formic acid in water and 0.1% in Acetonitrile. The LC system was coupled online to an orbitrap Eclipse™ Tribrid™ Mass Spectrometer (ThermoFisher Scientific) via an EasySpray ion source connected to a FAIMSPro device. The scan sequence began with an MS1 spectrum (Orbitrap analysis, resolution 120,000, scan range 400–1000 Th, automatic gain control (AGC) target of 300%, maximum IIT 50 ms, RF lens 40%). Sequential MS2s were collected from 500 to 900 Th with a scan range from 200 - 1200 Th. Each MS2 scan used higher-energy collisional dissociation (HCD) for fragmentation at a NCE (normalized collision energy) setting of 33%, and an AGC (automatic gain control) target set to 1000%. The nanospray ionization voltage was set to 2300 volts, FAIMSPro gas flow was set to static (3.6 L/min) and the temperature of the ion transfer tube was set to 240 °C. FAIMSPro ion mobility was applied (standard resolution) and its compensation voltage was set to −45 volts. The isolation window was 10 m/z for all benchmarking experiments except for the comparison of the windowing scheme. Resolution and maximum IIT were set for each experiment as described in the supplementary (Supporting information: Acquisition Parameters for each method).

### Data analysis

#### Raw data analysis and downstream data analysis

AlphaPept (version 0.4.5)^27^ was used to analyse DDA data for tryptic HeLa quality control. Spectronaut (version 15.5.211111.50606 and 16.0.220606.53000) (Biognosys)^28^ was used for raw data quantification and raw data were searched against the 9606 *homo sapiens* database obtained from Uniprot (Swiss-Prot with isoforms was downloaded on 07/11/2020) in a directDIA™ analysis. False-discovery rates (FDR) were set at 1% on peptide spectral match (PSM), peptide, and protein levels. Enzyme specificity was set to trypsin/P and lysC. The maximum allowed peptide length in search space was set to fifty-two and the minimum was set to seven. The maximum allowed number of missed cleavages in search space was set to two. Cysteine carbamidomethyl was set as a fixed modification. N-terminal acetylation and methionine oxidation were set as variable modifications. Default settings were applied for other parameters.

Spectronaut output tables were imported into RStudio (Version 2021.09.2+382) for proteomics data analysis and visualization. In this study, results were analysed based on the number of identified peptides. Briefly, peptides found in less than 65% of replicated experiments were removed from the analysis. Log transformation was performed on the peptide level to visualize its distribution between mass analyzers at various concentrations. Lastly, identified peptides from each method at various concentrations were shown as stacked bar charts with CV ranges between < 10%, 10-15%, and > 15%.

## RESULTS AND DISCUSSION

### Comparison between an orbitrap mass analyser and linear ion trap mass analyser on low-input proteomics

To evaluate the impact of using a LIT instead of an OT for MS2 readouts in DIA, we compared the two mass analyzers on an Eclipse™ Tribrid™ Mass Spectrometer (ThermoFisher Scientific). We focused especially on the performance of these mass analyzers on low-input samples, by measuring a dilution series ranging from 100 ng down to 1 ng. In line with previous results^19^, we find that LIT-DIA starts to outperform OT-DIA on samples below 10 ng load (Fig. 1A). Conversely, above 10 ng loads, the number of identified peptides from LIT-DIA does not improve substantially, while the number of identified peptides from OT-DIA proportionally increases with the increasing concentration of input material. This is likely due to the lower specificity of the LIT, hindering the deconvolution of very complex DIA spectra. Interestingly, in our setup we were able to identify 5x more peptides at 1 ng than Borras et al.^19^, using a 4x shorter gradient. Furthermore, compared to the result from using an orbitrap mass analyzer in Bekker-Jensen *et al*^23^, we could identify > 10,0000 precursors and > 2,300 protein groups with 40 SPD 31 minutes LC gradients using LIT-DIA-FAIMS from 5 ng of a HeLa digest.

**Fig. 1.**
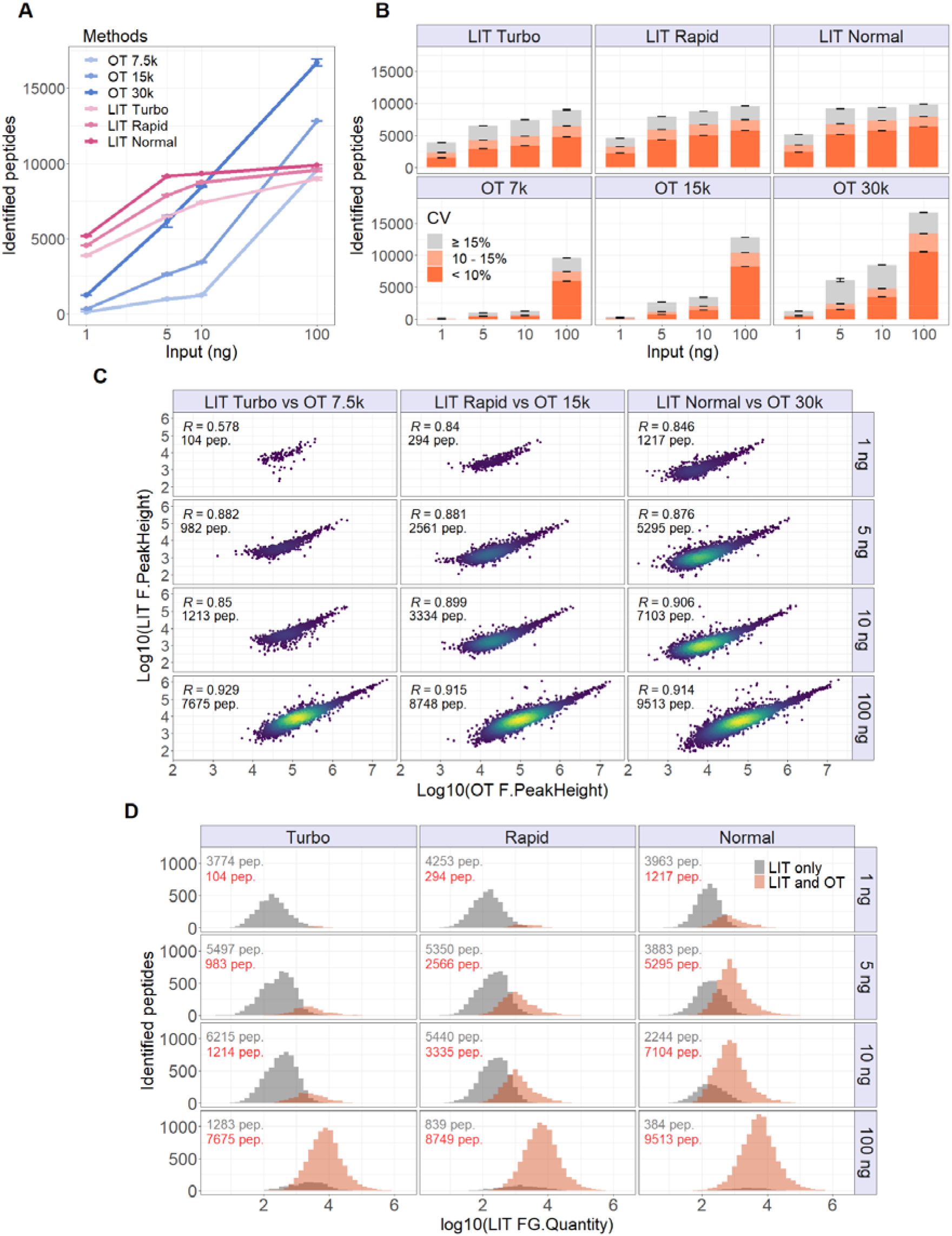
Comparison between an Orbitrap Mass analyzer and Linear Ion Trap Mass Analyzer on low-input proteomics. **A.** Comparison of the number of identified peptides in serial dilution (1, 5, 10, and 100 ng) of HeLa tryptic digest between OT-DIA-based methods with different resolution scans (7.5k, 15k, and 30k) and LIT-DIA-based methods with different scanning modes (*Turbo, Rapid, Normal* on 40 SPD). **B.** Comparison of identified peptides between the OT-DIA-based methods and the LIT-DIA-based methods. Identified peptides with a coefficient of variation (CV) between 10% and 15% are coloured with light red and those with a CV below 10% with dark red. **C.** Pearson correlation of fragment ion intensities between OT-DIA-based methods and LIT-DIA-based methods when 1, 5, 10, and 100 ng of HeLa tryptic digested were analyzed in quadruplicate. **D.** Distribution of the range of detection between the OT-DIA-based method and LIT-DIA-based methods and overlapping of identified peptides based on their intensities.

To investigate the precision of peptide quantification across replicate measurements, we calculated the coefficient of variation (CV) between technical replicates (Fig. 1B). We find that a high fraction of peptides has CVs below 10% for all methods. Interestingly, this fraction decreases in OT-DIA 30k, likely due to the longer cycle time (Table 2), resulting in fewer points per peak (PPP) and thus unreliable peak area estimation. Subsequently, we used the measured cycle times for each method (Table 2) to match methods with comparable cycle times and calculated Pearson correlations of peptide abundances between the pairs of OT- and LIT-based methods (Fig. 1C). We find that peptide abundances are well correlated, indicating good reproducibility between OT and LIT quantification, except for the comparison between LIT-DIA *Turbo* and OT-DIA 7.5k on 1 ng input material, where due to the limited sensitivity of OT at such low IITs, only 104 peptides were detected in OT. Similarly, we examined the overlap between identified peptides in OT-DIA and LIT-DIA with increasing input material (Fig. 1D). The results show that for lower inputs, peptides measured by both LIT and OT tend to be higher abundant than peptides measured by LIT only, further supporting the higher sensitivity of the LIT. Conversely, at 100 ng input material, almost all peptides that were identified in LIT-DIA were also identified in OT-DIA.

**Table 1.** The LIT acquisition rates from this study are provided below, as their terms are Thermo Scientific nomenclature.

| LIT Scanning Mode | Injection Time (ms) | Scanning Speed (Hz) | Time per Scan (ms) |
| --- | --- | --- | --- |
| Turbo | 16 | 38.5 | 26 |
| Rapid | 23 | 30.3 | 33 |
| Normal | 38 | 20.8 | 48 |
| Enhanced | 118 | 7.9 | 126 |
| Zoom | 452 | 1.9 | 522 |

**Table 2.** Parameters for method comparison between LIT-DIA-based methods and OT-DIA-based methods on 40 SPD LC method.

| Mass Analyzer | Scanning Mode | Injection Time (ms) | Cycle Time (s) | Windowing Scheme (m/z) | Average PPP |  |  |  |
| --- | --- | --- | --- | --- | --- | --- | --- | --- |
|  |  |  |  |  | 1 ng | 5 ng | 10 ng | 100 ng |
| Linear ion trap | Turbo | 16 | 1.4 | 10 | 7 | 7 | 7 | 15 |
| Linear ion trap | Rapid | 23 | 1.7 | 10 | 6 | 7 | 7 | 14 |
| Linear ion trap | Normal | 38 | 2.4 | 10 | 5 | 6 | 6 | 11 |

| Mass Analyzer | Resolution | Injection Time (ms) | Cycle Time (s) | Windowing Scheme (m/z) | Average PPP |  |  |  |
| --- | --- | --- | --- | --- | --- | --- | --- | --- |
|  |  |  |  |  | 1 ng | 5 ng | 10 ng | 100 ng |
| Orbitrap | 7500 | 11 | 1.5 | 10 | 8 | 7 | 6 | 10 |
| Orbitrap | 15000 | 22 | 1.9 | 10 | 5 | 5 | 5 | 9 |
| Orbitrap | 30000 | 54 | 3.2 | 10 | 4 | 4 | 4 | 6 |

To investigate the quantitative accuracy of peptide quantification across replicate measurements of LIT-DIA, we first quantified the reproducibility from 2 technical replicates from OT-DIA-based methods with different resolution scans (7.5k, 15k, and 30k) and LIT-DIA-based methods with different scanning modes (*Turbo, Rapid, Normal* on 40 SPD). at 1-, 5-, 10-, and 100 ng of HeLa tryptic digest, and the technical reproducibility of our measurements is satisfying with Pearson’s R = 0.9 on LIT in low-input (1-, 5-, and 10 ng) and 100 ng (Fig. 2A). For the low-input analysis, our results show that LIT (Pearson’s R = 0.9) is more precise than OT (Pearson’s R = 0.8) at the same scanning speed. Next, we quantified peptides from dilution series measurement to determine the quantitative accuracy of LIT-DIA compared to OT-DIA. Data were log-transformed and normalized so the median values of 1 ng were set to zero. Our results show that quantified peptides from both mass analyzers in the dilution series were linearly correlated with dilution factors as expected (Fig. 2B).

**Fig. 2.**
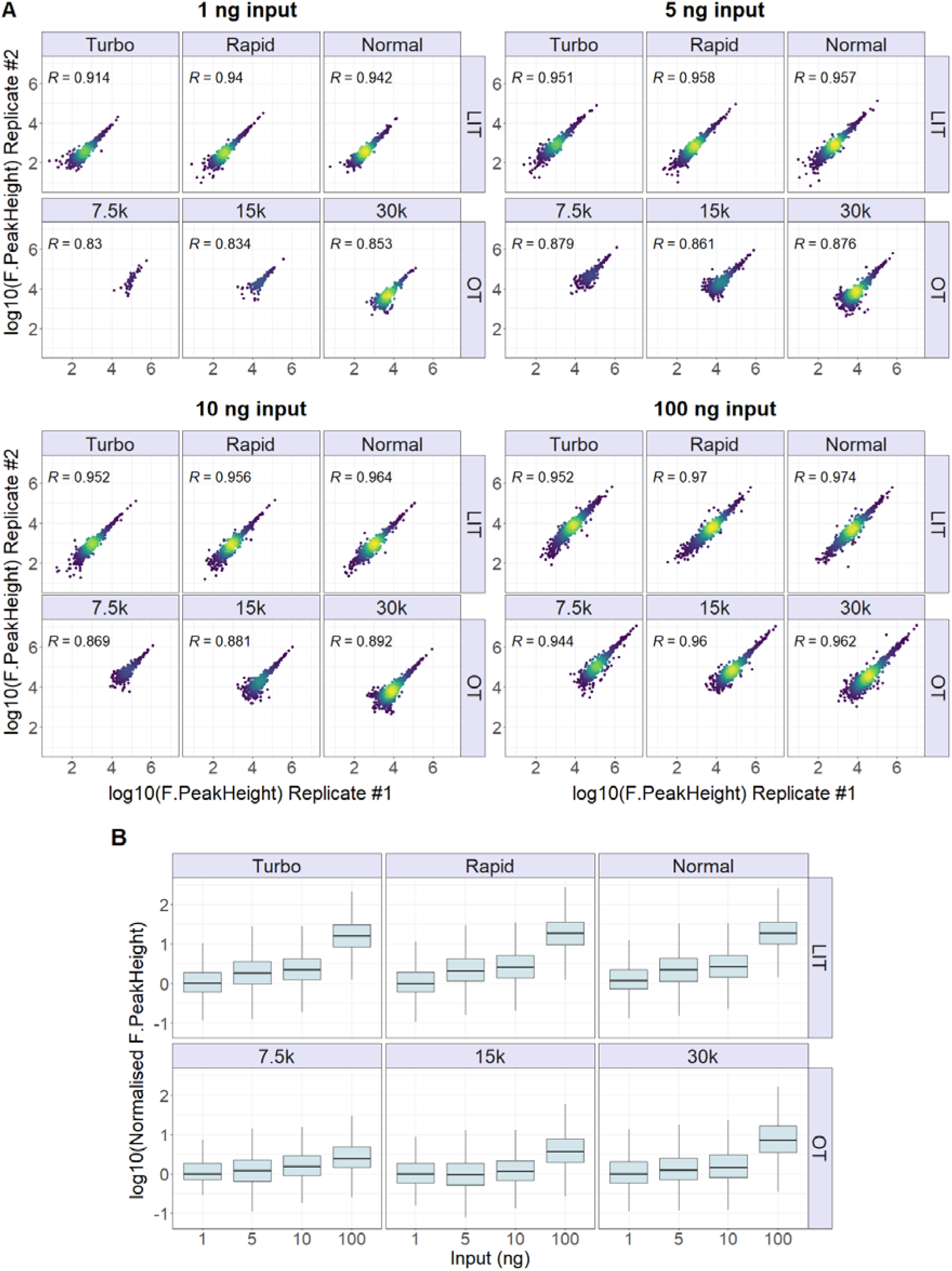
**A.** Quantitative reproducibility on peptide level of dilution series (1-, 5-, 10-, and 100 ng) of tryptic HeLa from technical replicates. **B.** The linear quantitative response curve on peptide level of the dilution series of tryptic HeLa (1-, 5-, 10-, and 100 ng).

### Exploring wide window data-independent acquisition for low-input proteomics

In order to determine the more optimal use of OT technology in low-input proteomics, we also investigated how isolation window widths affect peptide and protein identifications on OT-DIA and compared them with 10 m/z isolation windows on ‘*Norma*’ scanning mode with LIT-DIA. In low-input DIA, it becomes challenging to design an OT method to detect peptides at 10 m/z isolation windows, while keeping IITs high and PPP at acceptable levels. In other words, to successfully utilize an OT mass analyzer for sample inputs of 10ng and below, it is inevitable to widen the isolation windows and increase IITs in order to gain the required sensitivity. Conversely, due to increased IITs, we can also increase OT resolution to better match the OT transients with such longer IITs. Increasing the resolution on OT also tends to increase the signal-to-noise ratio of peaks due to increased ion oscillation during the longer transients, and thus induce a stronger ion current over time. This improvement was elegantly shown by Derks et al. in an approach termed plexDIA^15^, and therefore we analysed our HeLa dilution series with multiple OT-DIA methods with fixed cycle times, including the plexDIA method as a benchmark with Whisper™ 40 SPD LC method. The utilized acquisition schemes are described in Table 3 and raw data were analysed by Spectronaut version 16. Our results are summarized in Fig. 3, and it is evident that the number of identified peptides increases by means of increased IITs, with a concomitant increase in OT resolution and isolation window width. Especially for 1ng, the plexDIA method compares favourably to LIT-DIA, indicating the strength of wide-window OT-DIA for ultra-low loads as an alternative to narrow-window LIT-DIA.

**Fig. 3.**
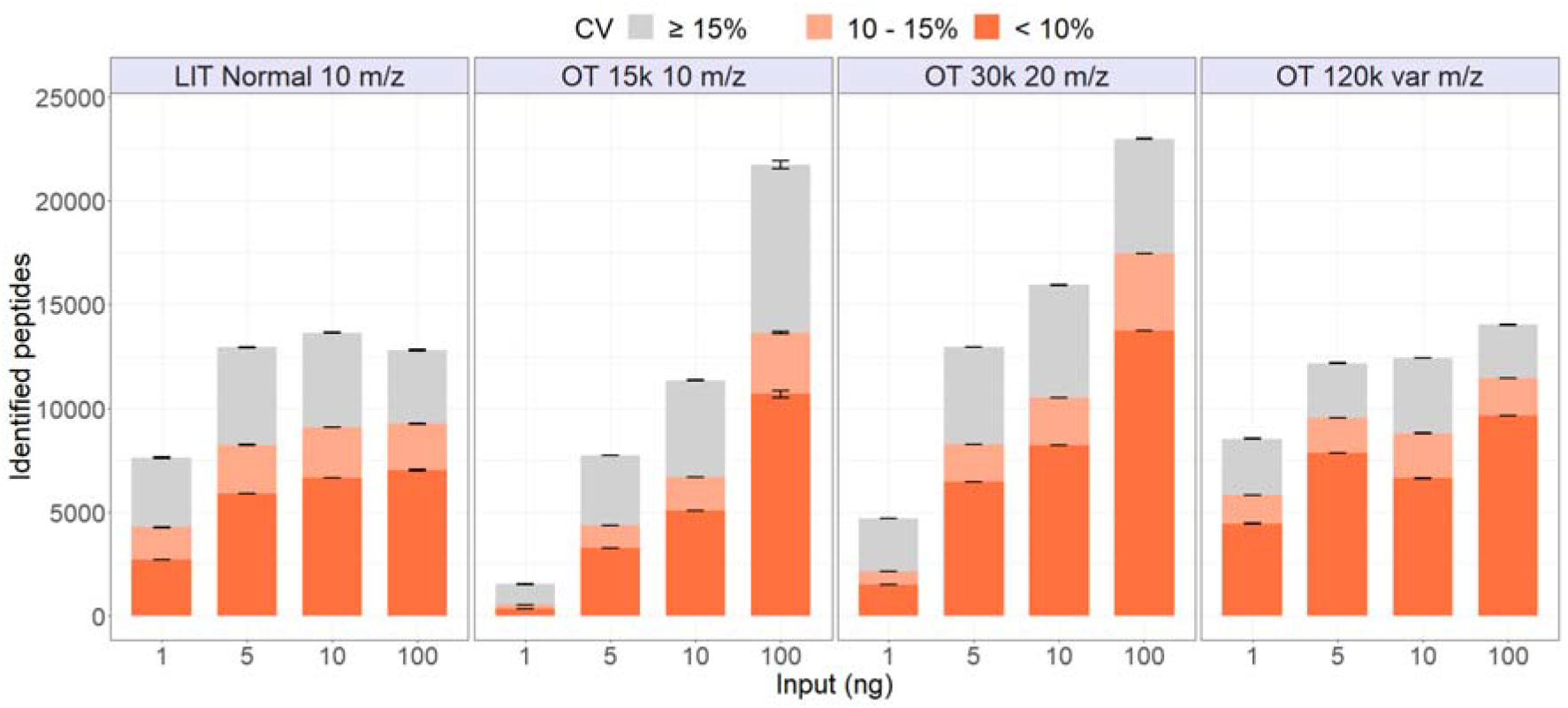
Exploring the wide window of data-independent acquisition for low-input proteomics. Comparison between LIT-DIA with 10 m/z isolation windows and OT-DIA methods (10 m/z isolation windows on 15k resolution, 20 m/z isolation windows on 30k resolution and various windows sizes (4 windows with 120-, 120-, 200-, and 580 m/z) on 120k resolution)

**Table 3.** Parameters for method comparison between LIT-DIA-based methods and OT-DIA-based methods on 40 SPD LC methods with wide isolation windows.

| Mass Analyzer | Scanning Mode | Injection Time (ms) | Windowing Scheme (m/z) | Number of Windows | Scan Range (m/z) |
| --- | --- | --- | --- | --- | --- |
| Linear ion trap | Normal | 38 | 10 | 40 | 500-900 |
| Mass Analyzer | Resolution | Injection Time (ms) | Windowing Scheme (m/z) | Number of Windows | Scan Range (m/z) |
| Orbitrap | 15000 | 22 | 10 | 40 | 500-900 |
| Orbitrap | 30000 | 54 | 20 | 20 | 500-900 |
| Orbitrap | 120000 | 246 | 120, 120, 200 and 580 | 4 | 378-1402 |

### Balancing an optimised windowing scheme with ion injection times

After testing the different mass analyzers, we set out to evaluate the impact of varying windowing schemes. We focused on improving peptide identification for low-input proteomics, and reproducibility of the workflow while maintaining acceptable cycle times. We tested LIT-DIA methods with 34, 40, and 45 windows with constant IITs set to auto (38 ms for LIT *Normal* scanning mode) covering a scan range of 500-900 m/z. We found that the windowing scheme reported previously^19^ also identified and quantified the highest number of peptides (Fig. 4). Based on our results, we opted to use the 40 windows scheme as a standard setup for all the evaluations in our study.

**Fig. 4.**
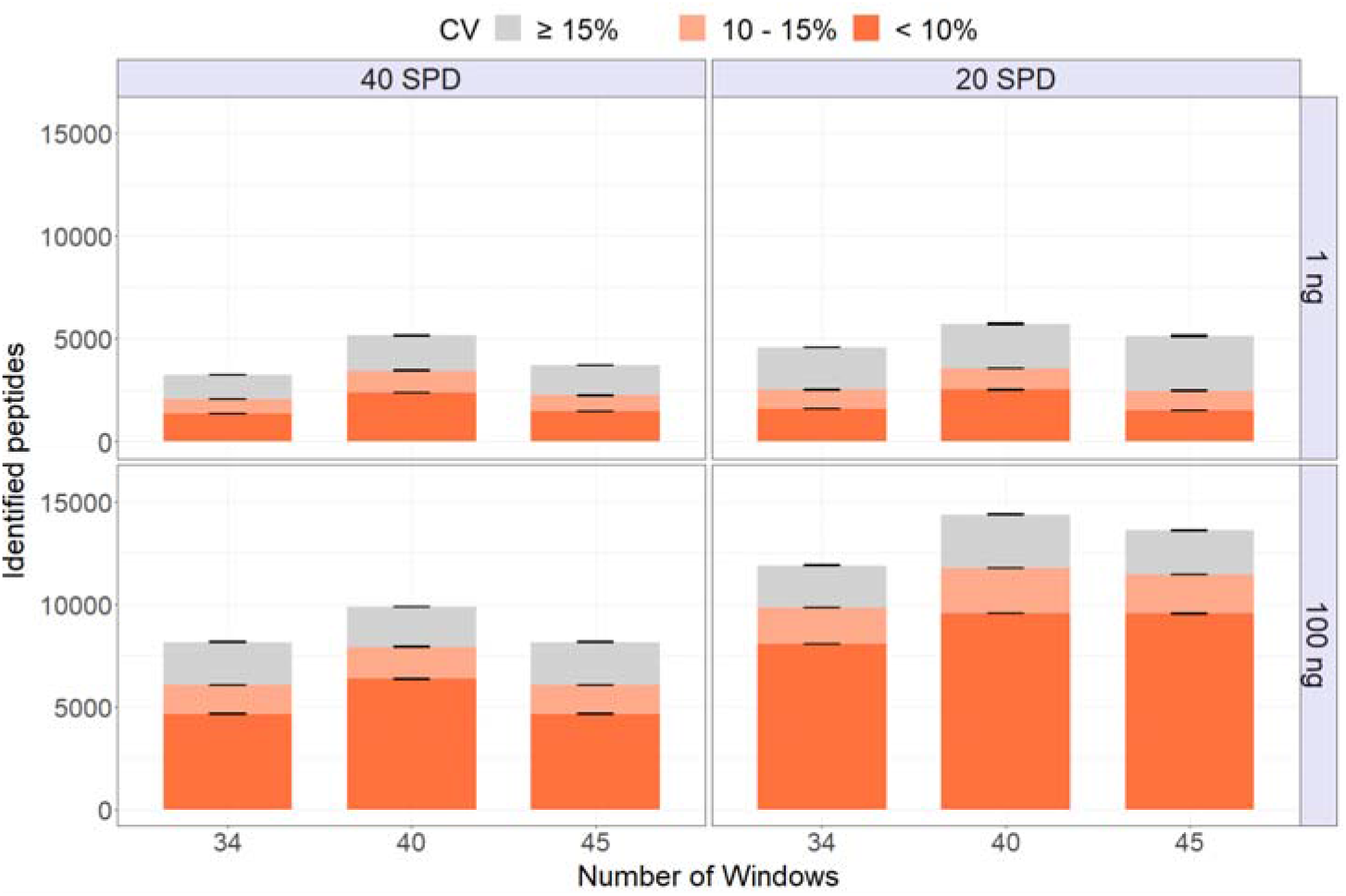
Comparison of windowing scheme. Comparison of the number of identified peptides on LIT-DIA-based method on *Normal* scanning mode with constant auto-IIT and different numbers of windows (34, 40, and 45 windows) and input material (low, and high-input tryptic HeLa digest) on Whisper™ 100 for 40 SPD and 20 SPD. Identified peptides with a coefficient of variation (CV) between 10% and 15% are coloured with light red and those with a CV below 10% with dark red.

Next, we repeated experiments to test the effect of using higher ion injection times (IITs) at similar cycle times, which comes at the cost of fewer, larger isolation windows to cover the same m/z range during the fixed cycle time (Supporting information: Sup. Fig. 1). Interestingly, we find that this strategy resulted in fewer identified peptides for a 1 ng sample, indicating that the increasing complexity of MS2 spectra derived from wider isolation windows presents a major challenge for LIT-DIA. Nonetheless, increasing IITs can be desirable to increase the number of ions sampled for subsequent fragmentation. Thus, we subsequently tested the effect of using higher IITs, while using a constant 40 DIA windows, resulting in increased cycle times (Fig. 5). Up to 80 ms IIT, the results indicate not only a steady increase in peptide identification but also a quantitative precision of those identifications. However, this increase plateaus beyond 80 ms, likely due to increased cycle times affecting efficient sampling of both MS1 and MS2 spectra.

**Fig. 5.**
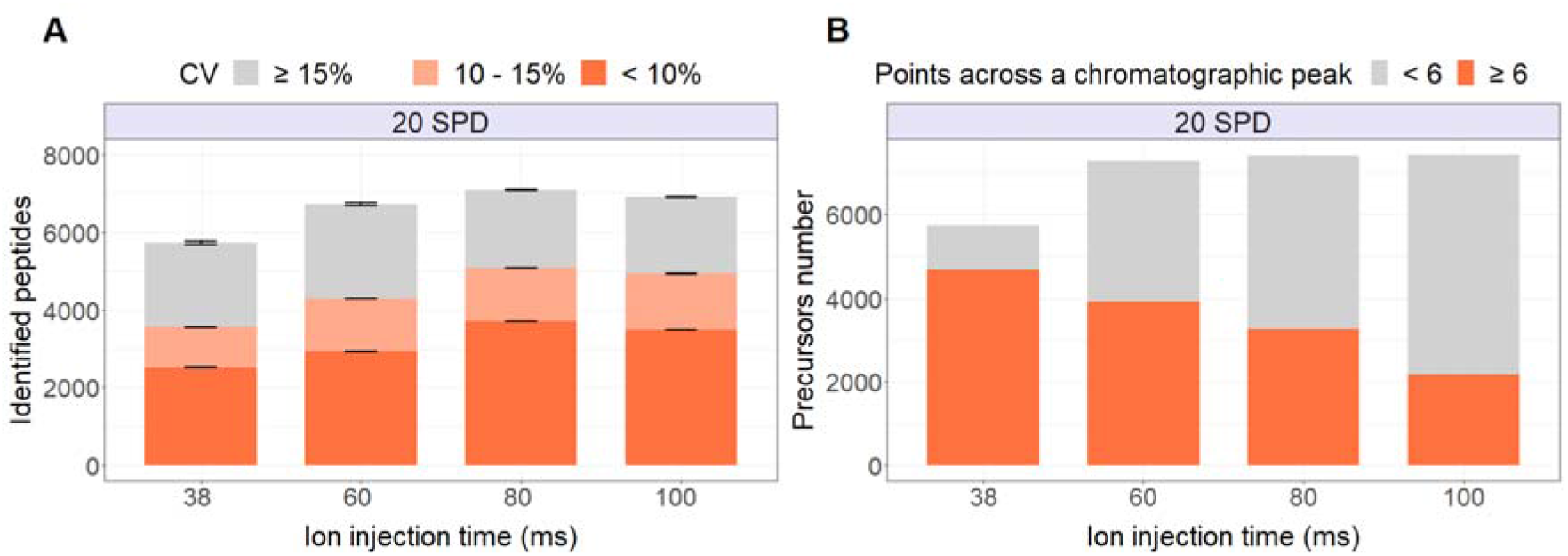
Increasing injection time on DIA LIT methods. **A.** Comparison of the number of identified peptides on LIT-DIA-based method on *Normal* scanning mode for Whisper™ 100 20 SPD from 1 ng of tryptic HeLa digest with different IITs at fixed 40 isolation windows. Identified peptides with a coefficient of variation (CV) between 10% and 15% are coloured with light red and those with a CV below 10% with dark red. The cycle times for the methods are indicated by their injection times: 38 ms 2.4 s, 60 ms 3.28 s, 80 ms 4.09 s, and 100 ms 4.91 s. **B.** Comparison of the number of precursors with a number of points across a peak equal to or greater than 6 on LIT-DIA-based method on *Normal* scanning mode from 1 ng input of tryptic HeLa digest on Whisper™ 100 20 SPD with different IITs (38 ms, 60 ms, 80 ms, and 100 ms) at fixed 40 isolation windows.

### Comparison of injection time on linear ion trap mass analyzer

To enhance the data quality for both identification and quantification, we tested LIT-DIA methods at different scanning modes and IITs. For a gradient flow of 100 nl/min with the 31 minutes method (Whisper™ 40 SPD), we evaluated LIT-DIA methods at *Turbo, Rapid*, and *Normal* scanning modes, and for a gradient flow of 100 nl/min with the 58 minutes method (Whisper™ 20 SPD), we evaluated LIT-DIA methods at *Rapid, Normal, Enhanced* and *Zoom* scanning modes. In all cases, IITs were set to auto to ensure optimal parallelization of ion accumulation and scan times in each scanning mode. We find that setting LIT-DIA-based methods with auto IIT and 40 isolation windows covering a scan range of 500-900 m/z is a good compromise between cycle time, reproducibility, and sensitivity for benchmarking low-input proteomics experiments based on our study. One exception is the LIT-DIA-based method using *Turbo* scanning mode, where besides the auto IIT (Fig. 6), we evaluated limiting IIT to 8 ms and found that its performance is similar to the LIT-DIA-based method on *Turbo* scanning mode at auto-IIT (16 ms) with 100 ng input-material. In contrast, with increasing IITs it is possible to identify more peptides but at a cost to quantitative precision due to increased cycle times.

**Fig. 6.**
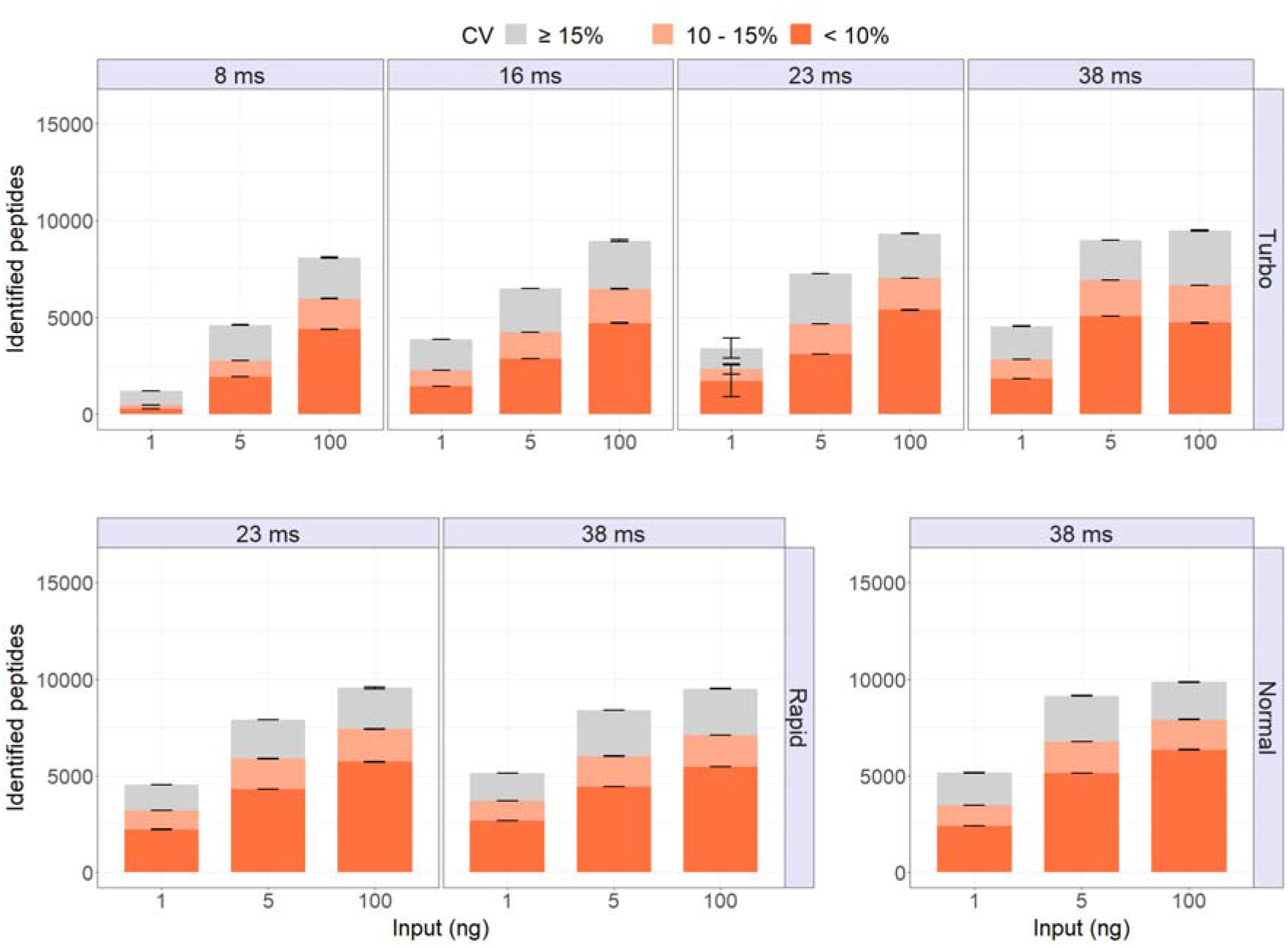
Comparison of Injection time on linear ion trap mass analyzer. Comparison of the number of identified peptides on LIT-DIA-based method on *Turbo, Rapid*, and *Normal* scanning mode for Whisper™ 100 40 SPD from 1,5, and 100 ng of tryptic HeLa digest with different IITs at fixed 40 isolation windows. For *Turbo* scanning mode auto IIT (16 ms), half of its auto-IIT (8 ms), and the auto IIT of *Rapid* (23 ms) and *Normal* (38 ms) were applied to determine the compromise between ion fill time and scanning speed of LIT on this mode. For *Rapid* scanning mode, the IITs of 23 ms and 38 ms suffice. For *Normal* scanning mode, only its auto-IIT was applied for this comparison. Identified peptides with a coefficient of variation (CV) between 10% and 15% are coloured with light red and those with a CV below 10% with dark red.

### Improvement of the number of points across chromatographic peaks

To reach a sufficient number of data points across chromatographic peaks for accurate quantification^29^, we tested different scanning modes (*Turbo, Rapid*, and *Normal* on Whisper™ 40 SPD and *Rapid, Normal*, and *Enhanced* on Whisper™ 20 SPD) to find a compromise between scanning mode and a number of data points across their chromatographic peak (i.e. points-per-peak, or PPP) for ultra-sensitive routine analysis.

For the low-input proteomics analysis with the LIT-DIA-based method, the *‘Rapid*’ scanning mode on 40 SPD appears to be the best fit because its Gaussian peaks have seven PPP as its median. On 20 SPD, the *‘Normal* scanning mode resulted in eight PPP as its median (Fig. 7A), suggesting in both cases that the quantification should be reliable^29^. Our results show that we can identify more peptides with a number of points across a peak lower than 6 by using the *“Normal”* scanning mode on 40 SPD, and the *“Enhanced”* scanning mode on 20 SPD (Fig. 7B). However, as visualized by the lower number of precursors with PPP > 6 at those scan rates, it is likely to affect the peak shape determination, leading to measurement errors.

**Fig. 7.**
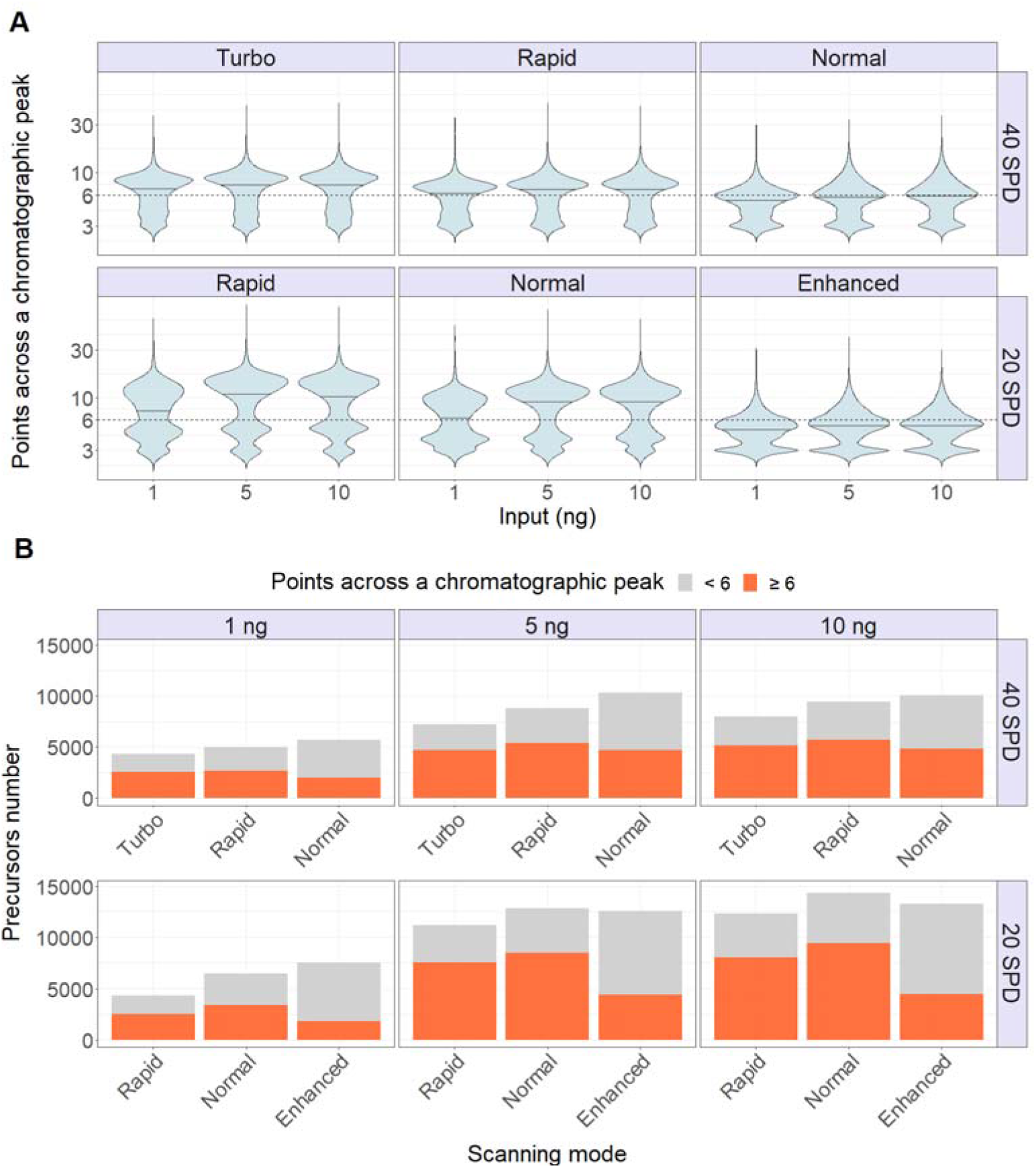
Improvement of the Number of Points Across Chromatographic Peaks. **A.** Distribution of numbers of points across a peak by using Whisper100™ 20 SPD with LIT-DIA-based methods on *Rapid, Normal*, and *Enhanced* scanning modes and Whisper100™ 40 SPD with LIT-DIA-based methods on *Turbo, Rapid*, and *Normal* scanning modes on 1, 5, and 10 ng input material. **B.** Comparison of the number of precursors with a number of points across a peak equal to or greater than 6 on the LIT-DIA-based method on *Turbo, Rapid*, and *Normal* scanning modes with different input material (1 ng, 5 ng, and 10 ng of tryptic HeLa digest) on Whisper™ 100 40 SPD and on *Rapid, Normal*, and *Enhanced* scanning mode with different input material (1 ng, 5 ng, and 10 ng of tryptic HeLa digest) on Whisper™ 100 20 SPD.

### Current Limitations

One of the bottlenecks for low-input proteomics experiments is that we do not have sufficient material to perform e.g. high-pH offline fractionation^30^ to reduce the complexity of biological samples, or gas-phase fractionation to efficiently generate a spectral library^31^. However, our results demonstrate that library-free quantification can establish a benchmark experiment on a complex tryptic digest. We expect that the number of identified proteins and peptides could be further improved by searching against spectral libraries to increase the identification performance and quantification precision of the library-free quantification. Compared to an OT, LIT mass analyzers have the superior sensitivity needed for low-input experiments. However, a tradeoff of using LIT is its noise level, affecting subsequent raw data analysis. Nevertheless, Liu et al^32^ have shown that it is possible to apply LIT for 6-plex TMT quantification with a LIT-HCD MS^3^ scan. Future software development that is tailor-made for LIT-DIA data analysis is required to interpret data produced by this acquisition approach fully. For the quantification in figure 2, samples were diluted in buffer A so it is possible that we were integrating background noise because the background is dropping at the same rate as the signal. Finally, as we are using very short gradients in this work (31 min and 58min methods run-to-run), peptide and protein identifications can be significantly increased by lengthening the runtime as previously tested by others^16,17^. Similarly, we opted not to include library-based searches or co-searching strategies with higher-load samples, not to confound our technical results by those enhancements obtained by search strategies offering ID transfer.

## CONCLUSIONS

This manuscript describes a DIA method tailored toward ultra-low-input sample analysis, relying on the combination of an OT mass analyzer for MS1 scans and a LIT mass analyzer for MS2. To reduce background contamination, we include a FAIMSPro interface and demonstrate the ability to quantify representative proteomes from minimal input material. Our results demonstrate the LIT mass analyzer is a powerful detector for low-input material proteomics for loads of 10 ng and below, and is well-suited for DIA-based analysis. We envision LIT-DIA to be an alternative way of analysing limited material samples, while still achieving the 10 m/z precursor isolation windows required for differentiating many sequence variants and PTMs (e.g.methionine oxidation) from canonical peptides.

Through a series of optimization steps, we find that the sensitivity of a LIT allows us to use very short IITs. While the slightly longer IITs used in conjunction with higher resolution LIT scans (e.g. *“Normal”* or *“Enhanced”*) produced slightly more protein IDs, the quantification quality of those proteins was hampered. We attribute this to the longer cycle times required for higher resolution LIT scans, thereby reducing the points across a chromatographic peak for quantification.

This study compares proteome depth by using pre-defined LC methods on the Evosep ONE instrument (20 SPD and 40 SPD), which is the persistent tradeoff between sample throughput and proteome depth. With the 20 SPD chromatography method, more peptides can be identified and quantified than in the 40 SPD method, but the proteome and peptide coverage does not scale linearly. Hence, it depends on the biological question at hand which method should be preferred, and the data from our analyses can help guide such decisions. Our data suggest the main advantage of LIT over OT mass analyzers is their higher sensitivity at similar IITs on low-load samples. We envision their extreme scan speeds, resulting in shorter cycle times, are potentially well-suited for higher sample loads. However, our comparison at 100 ng suggests that additional acquisition advancements such as e.g. BoxCar and MSX acquisitions or multi-CV FAIMS DIA^23,33,34^ will be required to offset the lower LIT resolution.

In this study, we used Spectronaut^TM^ software to analyze raw data in directDIA™ mode due to its reliability and robustness. At the onset of our experimental evaluations, we analyzed our data with Spectronaut™ version 15. With the recent release of version 16, we decided to test the impact of new implementations and benchmark the performance of the new machine learning framework and Artificial Intelligence (AI)-based peak identification feature when deployed on LIT-DIA raw data. Raw files were reanalyzed with version 16 using identical parameters as used in version 15 to make results comparable. Spectronaut version 16 demonstrates clearly improved performance in terms of IDs, with an average of 20% improvement in our study (Fig. S7). These results underline the performance enhancements that can be gained when a computational interpretation of complex spectra is improved through advanced machine learning.

Furthermore, we demonstrate that the Evosep ONE^35^ can increase the throughput and the sensitivity of low-input proteomics experiments. Future improvements to this method could include improved sample throughput by chemical multiplexing methods such as e.g. TMT labelling or Ac-IP tag^15,18,36–38^. Especially in the case of the latter, the additive signal effect on MS2 scans is likely to prove fruitful for ultra-low input applications such as laser capture microdissection (LCM) or scMS, thereby not only increasing sample throughput but also improving sensitivity and proteome depths.

We hope that this study will serve as a useful resource for low-input proteomics studies, inspire dedicated data processing improvement efforts, and provide relevant starting points for implementing LIT-DIA on compatible instrument platforms around the world.

## Supporting information

LIT-DIA Supplementary

## ASSOCIATED CONTENT

### Data availability

All mass spectrometry raw data and search engine files from Spectronaut versions 15 and 16 from this study have been deposited to the ProteomeXchange Consortium via the MassIVE repository. Project accession: PXD034862 (ftp://MSV000089718@massive.ucsd.edu)

All of the code used to generate and analyze MS output is available in the Schoof lab GitHub repository (https://github.com/Schoof-Lab/LITDIA).

### Supporting information

- Supporting figures: optimizing a balance between Windowing Schemes and IITs (Figure S1); Shared peptides and protein groups between 1 ng-, 5 ng-, 10 ng-, and 100 ng tryptic HeLa (Figure S2); Distribution of the intensities of peptides (left) or protein groups (right) from 100 ng of tryptic HeLa and overlapping of identified peptides and protein groups between 1 ng and 100 ng tryptic HeLa (Figure S3); Comparison of identified peptides between the OT-DIA-based methods and the LIT-DIA-based methods from 1 ng tryptic HeLa. Identified peptides with a coefficient of variation (CV) between 10% and 15% are coloured with light red and those with a CV below 10% with dark red (Figure S4); Proportions of peptides in the 3 CV groups (<10% in dark red, 10 - 15% in light red and >= 15% in grey) with LIT and OT mass analyzer at different scanning speeds (*Turbo, Rapid, Normal*), resolution (7000, 15000 and 3000) and inputs (1-, 5-, 10-, 100 ng) (Figure S5); Quantitative reproducibility on peptide level of 1-, and 100 ng tryptic HeLa from technical replicates (Figure S6); Comparison of low-input protein identifications between Spectronaut™ version 15 and 16. Comparison of the number of identified peptides in serial dilution (1,10, and 100 ng) of HeLa tryptic digested analyzed by LIT-DIA-based methods with *Normal* scanning mode on 40 SPD between Spectronaut version 15 and 16. (Figure S7).
- Supporting table S1-7: The number of identified peptides and protein groups in each method, its cycle time, and the version of the search engine (Spectronaut) are listed.
- Acquisition parameters for each method.

## AUTHOR INFORMATION

### Authors

**Teeradon Phlairaharn** - The Novo Nordisk Foundation Center for Protein Research, Faculty of Health Sciences, University of Copenhagen, Copenhagen, Denmark; Department of Proteomics and Signal Transduction, Max-Planck Institute of Biochemistry, Martinsried, Germany; Department of Chemistry, Technical University of Munich, Munich, Germany; Department of Biotechnology and Biomedicine, Technical University of Denmark, Lyngby, Denmark; Pelotonia Institute for Immuno-Oncology, The Ohio State University Comprehensive Cancer Center, Columbus, Ohio 43210, United States; Department of Biomedical Informatics, The Ohio State University, Columbus, Ohio 43210, United States;

**Samuel Grégoire** - Computational Biology Unit, de Duve Institute, Université Catholique de Louvain, Brussels, Belgium; Department of Biotechnology and Biomedicine, Technical University of Denmark, Lyngby, Denmark;

**Lukas R. Woltereck** - Department of Chemistry, Technical University of Munich, Munich, Germany; Department of Biotechnology and Biomedicine, Technical University of Denmark, Lyngby, Denmark;

**Valdemaras Petrosius** - Department of Biotechnology and Biomedicine, Technical University of Denmark, Lyngby, Denmark;

**Benjamin Furtwängler** - Department of Biotechnology and Biomedicine, Technical University of Denmark, Lyngby, Denmark; The Finsen Laboratory, Rigshospitalet, Faculty of Health Sciences, University of Copenhagen, Copenhagen, Denmark; Biotech Research and Innovation Center (BRIC), University of Copenhagen, Copenhagen, Denmark;

### Author Contributions

TP and EMS conceived and designed the project. TP, VP, and EMS performed the experiments. TP, BCS, and EMS performed method optimization. TP, SG, and BCS performed the data analysis and visualization. LRW and BF contributed with input to the method design and data evaluation. TP, BCS, and EMS drafted and revised the manuscript, which has been read and approved by all authors. BCS and EMS supervised the work.

### Notes

The authors declare no competing financial interests. BCS is a founder and shareholder in Proteome Software, which operates in the field of proteomics.

## ACKNOWLEDGMENTS

TP would like to acknowledge the financial support from M. Mann at the Department of Proteomics and Signal Transduction, Max-Planck Institute of Biochemistry. TP would like to thank J. V. Olsen, Z. Ye, and L. Niu at the Novo Nordisk Center for protein research and M. Wilhelm at the Technical University of Munich for fruitful discussions. We thank the DTU Proteomics Core for instrument support, for all members of the Cell Diversity Lab led by ES.

## ABBREVIATIONS

CV: compensation voltage
DDA: data-dependent acquisition
DIA: data-independent acquisition
FAIMS: high-field asymmetric waveform ion mobility spectrometry
IIT: ion injection time
LC: liquid chromatography
LIT: linear ion trap
OT: orbitrap
scMS: MS-based single-cell proteomics
SPD: samples per day
TMT: tandem mass tag

