## Supplementary material for "High sensitivity limited material proteomics empowered by data-independent acquisition on linear ion traps": LIT-DIA Supplementary

### Supplementary Materials

#### Supporting figure

##### Figure S1

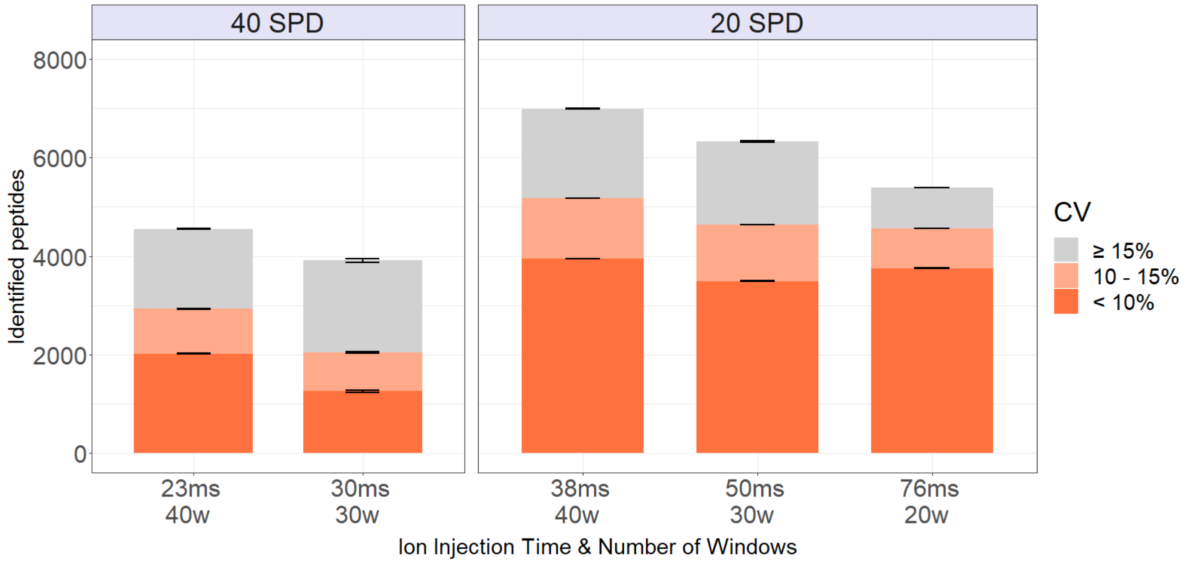


**Supporting figure S1 Optimizing a balance between Windowing Schemes and ITs.** Comparison of the number of identified peptides on DIA-LIT-based method on *Normal* scanning mode for Whisper™ 20 SPD and *Rapid* scanning mode for Whisper™ 40 SPD from 1 ng of tryptic HeLa lysate with different numbers of windows and IT at fixed cycle time. Identified peptides with a coefficient of variation (CV) between 10% and 15% are coloured with light red and those with a CV below 10% with dark red.

Figure S2

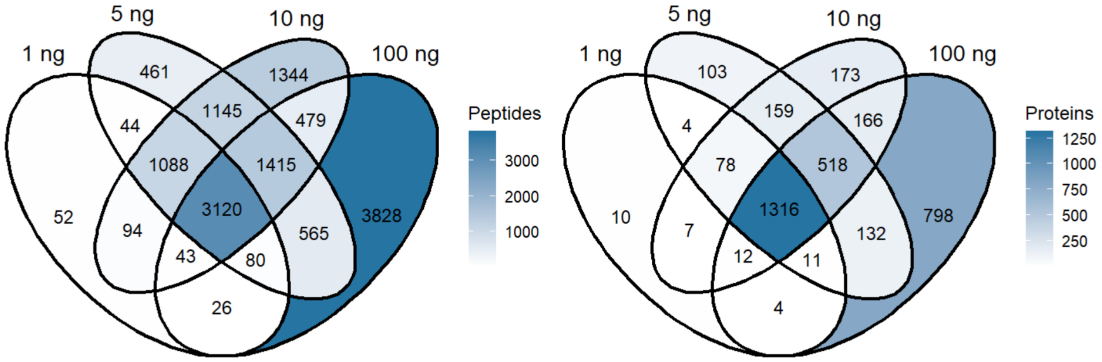


**Supporting figure S2 Shared peptides and protein groups between 1 ng-, 5 ng-, 10 ng-, and 100 ng tryptic HeLa.** A Venn diagram shows evidence that at low input material the LIT method still detects the same peptides as when used with higher amounts.

Figure S3

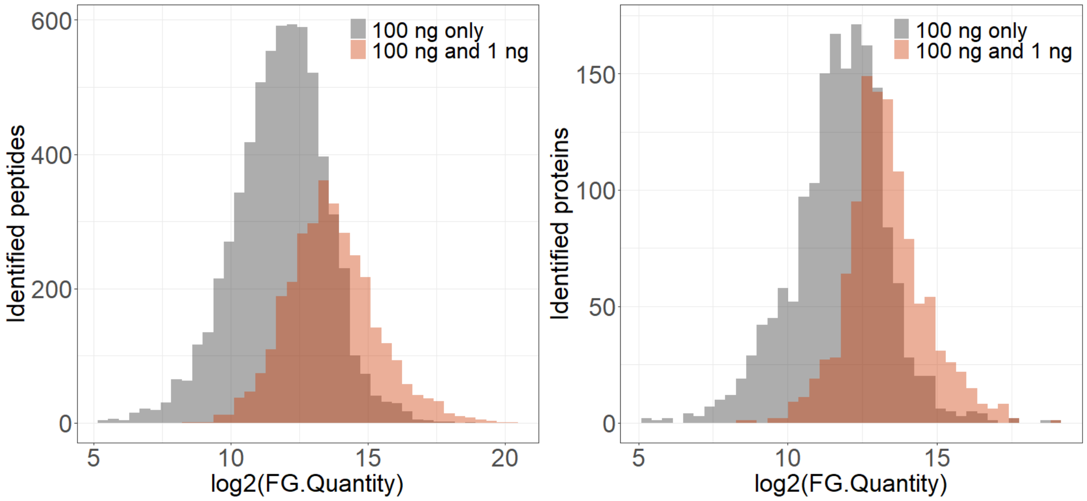


**Supporting figure S3 Distribution of the intensities of peptides (left) or protein groups (right) from 100 ng of tryptic HeLa and overlapping of identified peptides and protein groups between 1 ng and 100 ng tryptic HeLa.**

Figure S4

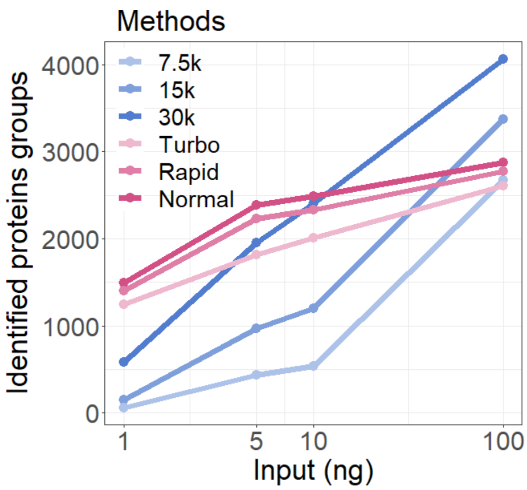


**Supporting figure S4 Comparison of the number of identified proteins groups in serial dilution (1, 5, 10, and 100 ng) of HeLa tryptic digested between DIA-OT-based methods with different resolution scans (7.5k, 15k, and 30k) and DIA-LIT-based methods with different scanning modes *(Turbo*, *Rapid*, *Normal* on 40 SPD) by Spectronaut version 15.**

Figure S5

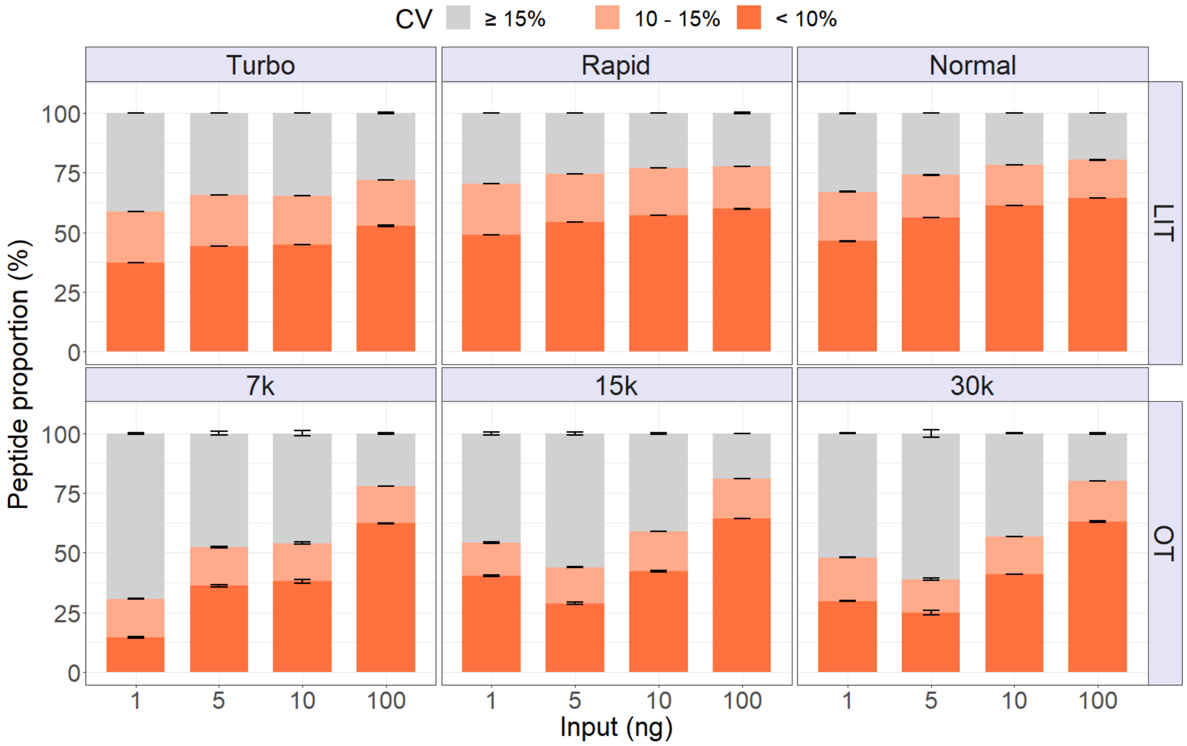


**Supporting figure S5** Proportions of peptides in the 3 CV groups (< 10% in dark red, between 10 and 15% in light red, and ≥ 15% in grey) with LIT and OT mass analyzer at different scanning speeds (*Turbo*, *Rapid*, *Normal*), resolution (7000, 15000 and 3000) and inputs (1-, 5-, 10-, 100 ng).

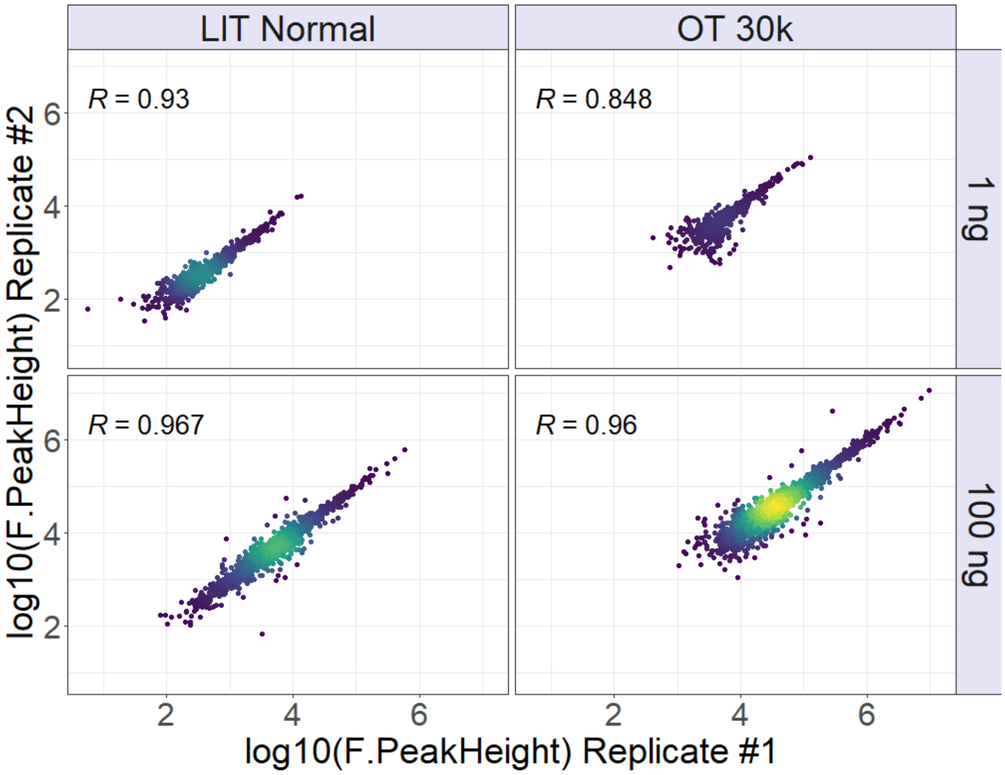


Figure S6

**Supporting figure S6 Quantitative reproducibility on peptide level of 1-, and 100 ng tryptic HeLa from technical replicates.**

Figure S7

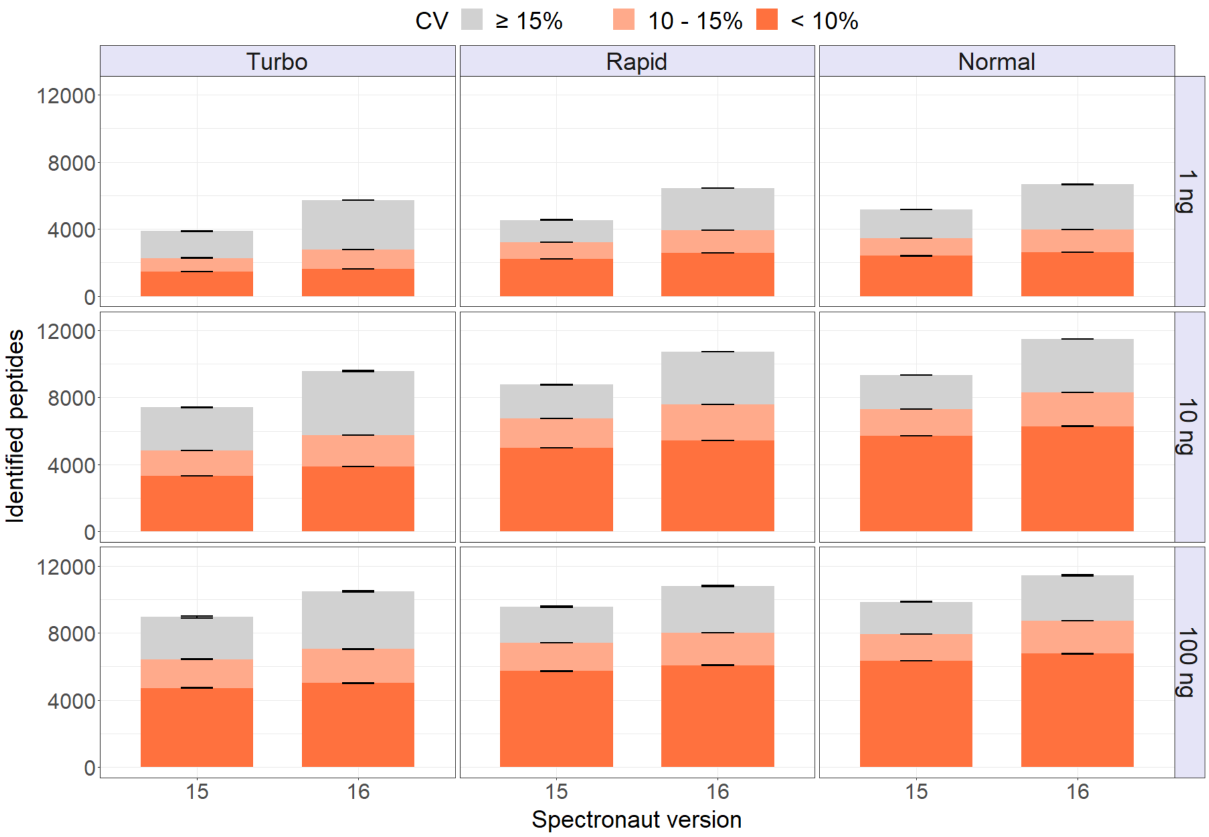


**Supporting figure S7 Comparison of low-input protein identifications between Spectronaut™ version 15 and 16. Comparison of the number of identified peptides in serial dilution (1,10, and 100 ng) of HeLa tryptic digested analyzed by DIA-LIT-based methods with *Normal* scanning mode on 40 SPD between Spectronaut version 15 and 16.**

#### Supporting tables

<u>Supporting table S1</u>

The number of identified peptides and protein groups in each method to compare the performance of both mass analyzers in different resolutions on OT and scanning modes on LIT at various concentrations of input material. Raw files were processed in Spectronaut version 15. The cycle time for each method was determined by Spectronaut version 15.

**Supplemental Table 1 for Fig. 1**

| Mass Analyser | Scanning Mode | Injection Time (ms) | Cycle Time (s) | Windowing Scheme (m/z) | Input (ng) | SPD | Spectronaut Version | Peptide ID | Protein Groups |
|---|---|---|---|---|---|---|---|---|---|
| LIT | Turbo | 16 | 1.46 | 10 | 1 | 40 | SN 15 | 3876 | 1243 |
| LIT | Turbo | 16 | 1.46 | 10 | 5 | 40 | SN 15 | 6465 | 1814 |
| LIT | Turbo | 16 | 1.47 | 10 | 10 | 40 | SN 15 | 7429 | 2013 |
| LIT | Turbo | 16 | 1.40 | 10 | 100 | 40 | SN 15 | 8976 | 2622 |
| LIT | Rapid | 23 | 1.75 | 10 | 1 | 40 | SN 15 | 4536 | 1402 |
| LIT | Rapid | 23 | 1.77 | 10 | 5 | 40 | SN 15 | 7886 | 2222 |
| LIT | Rapid | 23 | 1.77 | 10 | 10 | 40 | SN 15 | 8777 | 2335 |
| LIT | Rapid | 23 | 1.69 | 10 | 100 | 40 | SN 15 | 9561 | 2767 |
| LIT | Normal | 38 | 2.40 | 10 | 1 | 40 | SN 15 | 5160 | 1491 |
| LIT | Normal | 38 | 2.40 | 10 | 5 | 40 | SN 15 | 9160 | 2386 |
| LIT | Normal | 38 | 2.39 | 10 | 10 | 40 | SN 15 | 9331 | 2480 |
| LIT | Normal | 38 | 2.30 | 10 | 100 | 40 | SN 15 | 9886 | 2871 |
| **Mass Analyser** | **Resolution** | **Injection Time (ms)** | **Cycle Time (s)** | **Windowing Scheme (m/z)** | **Input (ng)** | **SPD** | **Spectronaut Version** | **Peptide ID** | **Protein Groups** |
| OT | 7500 | 11 | 1.52 | 10 | 1 | 40 | SN 15 | 106 | 62 |
| OT | 7500 | 11 | 1.52 | 10 | 5 | 40 | SN 15 | 1010 | 439 |
| OT | 7500 | 11 | 1.52 | 10 | 10 | 40 | SN 15 | 1275 | 541 |
| OT | 7500 | 11 | 1.90 | 10 | 100 | 40 | SN 15 | 9659 | 2684 |
| OT | 15000 | 22 | 1.96 | 10 | 1 | 40 | SN 15 | 300 | 150 |
| OT | 15000 | 22 | 1.96 | 10 | 5 | 40 | SN 15 | 2711 | 990 |
| OT | 15000 | 22 | 1.96 | 10 | 10 | 40 | SN 15 | 3466 | 1207 |
| OT | 15000 | 22 | 1.96 | 10 | 100 | 40 | SN 15 | 12871 | 3379 |
| OT | 30000 | 54 | 3.27 | 10 | 1 | 40 | SN 15 | 1267 | 591 |
| OT | 30000 | 54 | 3.27 | 10 | 5 | 40 | SN 15 | 6612 | 2038 |
| OT | 30000 | 54 | 3.27 | 10 | 10 | 40 | SN 15 | 8605 | 2432 |
| OT | 30000 | 54 | 3.21 | 10 | 100 | 40 | SN 15 | 16732 | 4062 |

Supporting table S2

The number of identified peptides and protein groups in each method to compare the performance of LIT on different numbers and sizes of isolation windows with 1 ng of HeLa tryptic lysate. For both LC methods (20-, and 40 SPD), scanning mode was set to *Normal*. Raw files were processed in Spectronaut version 15. The cycle time for each method was determined by Spectronaut version 15.

**Supplemental Table 2 for Fig. 2**

| Mass Analyser | Scanning Mode | Injection Time (ms) | Cycle Time (s) | Windowing Scheme (m/z) | Input (ng) | SPD | Spectronaut Version | Peptide ID | Protein Groups |
|---|---|---|---|---|---|---|---|---|---|
| LIT | Normal | 38 | 2.10 | 11.8 | 1 | 40 | SN 15 | 3239 | 1056 |
| LIT | Normal | 38 | 2.04 | 11.8 | 100 | 40 | SN 15 | 8127 | 2482 |
| LIT | Normal | 38 | 2.40 | 10 | 1 | 40 | SN 15 | 5160 | 1491 |
| LIT | Normal | 38 | 2.30 | 10 | 100 | 40 | SN 15 | 9886 | 2871 |
| LIT | Normal | 38 | 2.64 | 8.9 | 1 | 40 | SN 15 | 3734 | 1249 |
| LIT | Normal | 38 | 2.55 | 8.9 | 100 | 40 | SN 15 | 9675 | 2805 |
| LIT | Normal | 38 | 2.11 | 11.8 | 1 | 20 | SN 15 | 4568 | 1487 |
| LIT | Normal | 38 | 2.06 | 11.8 | 100 | 20 | SN 15 | 11978 | 3107 |
| LIT | Normal | 38 | 2.40 | 10 | 1 | 20 | SN 15 | 5736 | 1944 |
| LIT | Normal | 38 | 2.31 | 10 | 100 | 20 | SN 15 | 14020 | 3546 |
| LIT | Normal | 38 | 2.63 | 8.9 | 1 | 20 | SN 15 | 5773 | 1619 |
| LIT | Normal | 38 | 2.59 | 8.9 | 100 | 20 | SN 15 | 13519 | 3425 |

Supporting table S3

The number of identified peptides and protein groups in each method to compare the performance of LIT on different injection times and sizes of isolation windows (fixed cycle time) with 1 ng of HeLa tryptic lysate. The scanning mode for the 20 SPD LC method was set to *Normal* and *Rapid* for the 40 SPD LC method. Raw files were processed in Spectronaut version 15. The cycle time for each method was determined by Spectronaut version 15.

**Supplemental Table 3 for Fig. 3**

| Mass Analyser | Scanning Mode | Injection Time (ms) | Cycle Time (s) | Windowing Scheme (m/z) | Input (ng) | SPD | Spectronaut Version | Peptide ID | Protein Groups |
|---|---|---|---|---|---|---|---|---|---|
| LIT | Normal | 38 | 2.40 | 10 | 1 | 20 | SN 15 | 5773 | 1619 |
| LIT | Normal | 60 | 3.28 | 10 | 1 | 20 | SN 15 | 6819 | 1897 |
| LIT | Normal | 80 | 4.09 | 10 | 1 | 20 | SN 15 | 7123 | 2174 |
| LIT | Normal | 100 | 4.91 | 10 | 1 | 20 | SN 15 | 6940 | 2137 |

| Mass Analyser | Scanning Mode | Injection Time (ms) | Cycle Time (s) | Windowing Scheme (m/z) | Input (ng) | SPD | Spectronaut Version | Precorsus | Protein Groups |
|---|---|---|---|---|---|---|---|---|---|
| LIT | Normal | 38 | 2.40 | 10 | 1 | 20 | SN 15 | 6475 | 1619 |
| LIT | Normal | 60 | 3.28 | 10 | 1 | 20 | SN 15 | 7400 | 1897 |
| LIT | Normal | 80 | 4.09 | 10 | 1 | 20 | SN 15 | 7449 | 2174 |
| LIT | Normal | 100 | 4.91 | 10 | 1 | 20 | SN 15 | 7451 | 2137 |

<u>Supporting table S4</u>

The number of identified peptides and protein groups in each method to compare the performance of LIT on different injection times with 1 ng of HeLa tryptic lysate. The scanning mode was set to *Normal* coupled with the 20 SPD LC method. Raw files were processed in Spectronaut version 15. The cycle time for each method was determined by Spectronaut version 15.

**Supplemental Table 4 for Fig. 4**

| Mass Analyser | Scanning Mode | Injection Time (ms) | Cycle Time (s) | Windowing Scheme (m/z) | Input (ng) | SPD | Spectronaut Version | Peptide ID | Protein Groups |
|---|---|---|---|---|---|---|---|---|---|
| LIT | Turbo | 8 | 1.34 | 10 | 1 | 40 | SN 15 | 1216 | 515 |
| LIT | Turbo | 8 | 1.32 | 10 | 5 | 40 | SN 15 | 4621 | 1462 |
| LIT | Turbo | 8 | 1.29 | 10 | 100 | 40 | SN 15 | 8026 | 2314 |
| LIT | Turbo | 16 | 1.46 | 10 | 1 | 40 | SN 15 | 3876 | 1243 |
| LIT | Turbo | 16 | 1.46 | 10 | 5 | 40 | SN 15 | 6465 | 1814 |
| LIT | Turbo | 16 | 1.40 | 10 | 100 | 40 | SN 15 | 8976 | 2622 |
| LIT | Turbo | 23 | 1.76 | 10 | 1 | 40 | SN 15 | 4342 | 1334 |
| LIT | Turbo | 23 | 1.76 | 10 | 5 | 40 | SN 15 | 7238 | 2042 |
| LIT | Turbo | 23 | 1.6 | 10 | 100 | 40 | SN 15 | 9242 | 2658 |
| LIT | Turbo | 38 | 2.37 | 10 | 1 | 40 | SN 15 | 4549 | 1358 |
| LIT | Turbo | 38 | 2.37 | 10 | 5 | 40 | SN 15 | 8955 | 2297 |
| LIT | Turbo | 38 | 1.96 | 10 | 100 | 40 | SN 15 | 9558 | 2790 |
| LIT | Rapid | 23 | 1.75 | 10 | 1 | 40 | SN 15 | 4536 | 1402 |
| LIT | Rapid | 23 | 1.77 | 10 | 5 | 40 | SN 15 | 7886 | 2222 |
| LIT | Rapid | 23 | 1.69 | 10 | 100 | 40 | SN 15 | 9561 | 2767 |
| LIT | Rapid | 38 | 2.38 | 10 | 1 | 40 | SN 15 | 5147 | 1561 |
| LIT | Rapid | 38 | 2.38 | 10 | 5 | 40 | SN 15 | 8395 | 2306 |
| LIT | Rapid | 38 | 2.05 | 10 | 100 | 40 | SN 15 | 9527 | 2773 |
| LIT | Normal | 38 | 2.40 | 10 | 1 | 40 | SN 15 | 5160 | 1491 |
| LIT | Normal | 38 | 2.40 | 10 | 5 | 40 | SN 15 | 9160 | 2386 |
| LIT | Normal | 38 | 2.30 | 10 | 100 | 40 | SN 15 | 9886 | 2871 |

Supporting table S5

The number of identified peptides and protein groups in each method to compare the performance of LIT on different injection times and scanning modes with 1-,5-, and 100 ng of HeLa tryptic lysate on 40 SPD LC method. Raw files were processed in Spectronaut version 15. The cycle time for each method was determined by Spectronaut version 15.

**Supplemental Table 5 for Fig. 5**

| Mass Analyser | Scanning Mode | Injection Time (ms) | Cycle Time (s) | Windowing Scheme (m/z) | Input (ng) | SPD | Spectronaut Version | Precursors | Protein Groups |
|---|---|---|---|---|---|---|---|---|---|
| LIT | Turbo | 16 | 1.46 | 10 | 1 | 40 | SN 15 | 4291 | 1243 |
| LIT | Turbo | 16 | 1.46 | 10 | 5 | 40 | SN 15 | 7234 | 1814 |
| LIT | Turbo | 16 | 1.47 | 10 | 10 | 40 | SN 15 | 8012 | 2013 |
| LIT | Rapid | 23 | 1.75 | 10 | 1 | 40 | SN 15 | 4983 | 1402 |
| LIT | Rapid | 23 | 1.77 | 10 | 5 | 40 | SN 15 | 8829 | 2222 |
| LIT | Rapid | 23 | 1.77 | 10 | 10 | 40 | SN 15 | 9502 | 2335 |
| LIT | Normal | 38 | 2.40 | 10 | 1 | 40 | SN 15 | 5725 | 1491 |
| LIT | Normal | 38 | 2.40 | 10 | 5 | 40 | SN 15 | 10369 | 2836 |
| LIT | Normal | 38 | 2.39 | 10 | 10 | 40 | SN 15 | 10100 | 2480 |
| LIT | Rapid | 23 | 1.77 | 10 | 1 | 20 | SN 15 | 4348 | 1341 |
| LIT | Rapid | 23 | 1.77 | 10 | 5 | 20 | SN 15 | 11144 | 2689 |
| LIT | Rapid | 23 | 1.77 | 10 | 10 | 20 | SN 15 | 12352 | 2783 |
| LIT | Normal | 38 | 2.40 | 10 | 1 | 20 | SN 15 | 6475 | 1619 |
| LIT | Normal | 38 | 2.40 | 10 | 5 | 20 | SN 15 | 12775 | 3036 |
| LIT | Normal | 38 | 2.40 | 10 | 10 | 20 | SN 15 | 14309 | 3140 |
| LIT | Enhanced | 113 | 5.59 | 10 | 1 | 20 | SN 15 | 7567 | 1930 |
| LIT | Enhanced | 113 | 5.59 | 10 | 5 | 20 | SN 15 | 12642 | 3180 |
| LIT | Enhanced | 113 | 5.59 | 10 | 10 | 20 | SN 15 | 13106 | 3176 |

Supporting table S6

Raw files from LIT-DIA on different scanning modes with 1, 10, and 100 ng of HeLa tryptic lysate on the 40 SPD LC method were processed in Spectronaut versions 15 and 16 with identical settings to compare their performance on LIT-generated raw spectra. The number of identified peptides and protein groups in each method from both versions are listed.

**Supplemental Table 6 for Fig. 6**

| Mass Analyser | Scanning Mode | Injection Time (ms) | Cycle Time (s) | Windowing Scheme (m/z) | Input (ng) | SPD | Spectronaut Version | Peptide ID | Protein Groups |
|---|---|---|---|---|---|---|---|---|---|
| LIT | Turbo | 16 | 1.46 | 10 | 1 | 40 | SN 15 | 3876 | 1243 |
| LIT | Turbo | 16 | 1.47 | 10 | 10 | 40 | SN 15 | 7429 | 2013 |
| LIT | Turbo | 16 | 1.40 | 10 | 100 | 40 | SN 15 | 8976 | 2622 |
| LIT | Rapid | 23 | 1.75 | 10 | 1 | 40 | SN 15 | 4536 | 1402 |
| LIT | Rapid | 23 | 1.77 | 10 | 10 | 40 | SN 15 | 8777 | 2335 |
| LIT | Rapid | 23 | 1.69 | 10 | 100 | 40 | SN 15 | 9561 | 2767 |
| LIT | Normal | 38 | 2.40 | 10 | 1 | 40 | SN 15 | 5160 | 1491 |
| LIT | Normal | 38 | 2.39 | 10 | 10 | 40 | SN 15 | 9331 | 2480 |
| LIT | Normal | 38 | 2.30 | 10 | 100 | 40 | SN 15 | 9886 | 2871 |
| LIT | Turbo | 16 | 1.46 | 10 | 1 | 40 | SN 16 | 5716 | 1683 |
| LIT | Turbo | 16 | 1.47 | 10 | 10 | 40 | SN 16 | 9597 | 2505 |
| LIT | Turbo | 16 | 1.40 | 10 | 100 | 40 | SN 16 | 10467 | 2896 |
| LIT | Rapid | 23 | 1.76 | 10 | 1 | 40 | SN 16 | 6449 | 1908 |
| LIT | Rapid | 23 | 1.77 | 10 | 10 | 40 | SN 16 | 10703 | 2748 |
| LIT | Rapid | 23 | 1.68 | 10 | 100 | 40 | SN 16 | 10719 | 3008 |
| LIT | Normal | 38 | 2.40 | 10 | 1 | 40 | SN 16 | 6667 | 1825 |
| LIT | Normal | 38 | 2.39 | 10 | 10 | 40 | SN 16 | 11480 | 2867 |
| LIT | Normal | 38 | 2.30 | 10 | 100 | 40 | SN 16 | 11514 | 3125 |

Supporting table S7

The number of identified peptides and protein groups in each method to compare the performance of LIT on different injection times, numbers, and sizes of isolation windows with 1 ng and 100 ng of HeLa tryptic lysate. For both LC methods (20-, and 40 SPD), scanning mode was set to *Rapid* for 40 SPD and *Normal* for 20 SPD. Raw files were processed in Spectronaut version 15. The cycle time for each method was determined by Spectronaut version 15.

**Supplemental Table 7 for Sup. Fig. 1**

| Mass Analyser | Scanning Mode | Injection Time (ms) | Cycle Time (s) | Windowing Scheme (m/z) | Input (ng) | SPD | Spectronaut Version | Peptide ID | Protein Groups |
|---|---|---|---|---|---|---|---|---|---|
| LIT | Rapid | 23 | 1.75 | 10 | 1 | 40 | SN 15 | 4565 | 1405 |
| LIT | Rapid | 30 | 1.62 | 13.3 | 1 | 40 | SN 15 | 3973 | 1285 |
| LIT | Normal | 38 | 2.63 | 10 | 1 | 20 | SN 15 | 6994 | 1944 |
| LIT | Normal | 50 | 2.40 | 13.3 | 1 | 20 | SN 15 | 6312 | 1811 |
| LIT | Normal | 76 | 2.19 | 20 | 1 | 20 | SN 15 | 5394 | 1612 |

### Acquisition parameters for each method

#### DIA-OT 40X10 7.5K

The scan sequence began with an MS1 spectrum (Orbitrap analysis, resolution 120,000, scan range 400–1000 Th, automatic gain control (AGC) target of 300%, maximum injection time 50 ms, RF lens 40%). The precursor's mass range for MS2 analysis (Orbitrap analysis, resolution 7,500) was set from 500 to 900 Th and the scan range from 200 - 1200 Th. MS2 analysis consisted of higher-energy collisional dissociation (HCD), MS2 AGC was set to 1000%, NCE (normalized collision energy) was 33%, and the nanospray ionization was used in this experiment (positive ion 2300 volts, negative ion 600 volts, the gas mode was set to static, the temperature of ion transfer tube was set to 240 °C). FAIMSPro ion mobility was applied (Standard resolution, total carrier gas flow static at 3.6 liters per minute, and its compensation voltage was set to -45 volts). The isolation window was 10 Th over the mass range.

#### DIA-OT 40X10 15K

The scan sequence began with an MS1 spectrum (Orbitrap analysis, resolution 120,000, scan range 400–1000 Th, automatic gain control (AGC) target of 300%, maximum injection time 50 ms, RF lens 40%). The precursor's mass range for MS2 analysis (Orbitrap analysis, resolution 15,000) was set from 500 to 900 Th and the scan range from 200 - 1200 Th. MS2 analysis consisted of higher-energy collisional dissociation (HCD), MS2 AGC was set to 1000%, NCE (normalized collision energy) was 33%, and the nanospray ionization was used in this experiment (positive ion 2300 volts, negative ion 600 volts, the gas mode was set to static, the temperature of ion transfer tube was set to 240 °C).

FAIMSPro ion mobility was applied (Standard resolution, total carrier gas flow static at 3.6 liters per minute, and its compensation voltage was set to -45 volts). The isolation window was 10 Th over the mass range.

#### DIA-OT 40X10 30K

The scan sequence began with an MS1 spectrum (Orbitrap analysis, resolution 120,000, scan range 400–1000 Th, automatic gain control (AGC) target of 300%, maximum injection time 50 ms, RF lens 40%). The precursor's mass range for MS2 analysis (Orbitrap analysis, resolution 30,000) was set from 500 to 900 Th and the scan range from 200 - 1200 Th. MS2 analysis consisted of higher-energy collisional dissociation (HCD), MS2 AGC was set to 1000%, NCE (normalized collision energy) was 33%, and the nanospray ionization was used in this experiment (positive ion 2300 volts, negative ion 600 volts, the gas mode was set to static, the temperature of ion transfer tube was set to 240 °C). FAIMSPro ion mobility was applied (Standard resolution, total carrier gas flow static at 3.6 liters per minute, and its compensation voltage was set to -45 volts). The isolation window was 10 Th over the mass range.

#### DIA-OT 4Xvar 120K

The scan sequence began with an MS1 spectrum (Orbitrap analysis, resolution 120,000, scan range 400–1000 Th, automatic gain control (AGC) target of 300%, maximum injection time 50 ms, RF lens 40%). The precursor's mass range for MS2 analysis (Orbitrap analysis, resolution 120,000) was set from 378 to 1402 Th and the scan range for each scan from 200 - 1200 Th. MS2 analysis consisted of higher-energy collisional dissociation (HCD), MS2 AGC was set to 1000%, NCE (normalized collision energy) was 33%, and the nanospray ionization was used in this experiment (positive ion 2300 volts, negative ion 600 volts, the gas mode was set to static, the temperature of ion transfer tube was set to 240 °C). FAIMSPro ion mobility was applied (Standard resolution, total carrier gas flow static at 3.6 liters per minute, and its compensation voltage was set to -45 volts). The isolation windows were 120-, 120-, 200-, and 580 Th spread over the designated MS1 mass range.

#### DIA-OT 20X20 30K

The scan sequence began with an MS1 spectrum (Orbitrap analysis, resolution 120,000, scan range 400–1000 Th, automatic gain control (AGC) target of 300%, maximum injection time 50 ms, RF lens 40%). The precursor's mass range for MS2 analysis (Orbitrap analysis, resolution 30,000) was set from 500 to 900 Th and the scan range from 200 - 1200 Th. MS2 analysis consisted of higher-energy collisional dissociation (HCD), MS2 AGC was set to 1000%, NCE (normalized collision energy) was 33%, and the nanospray ionization was used in this experiment (positive ion 2300 volts, negative ion 600 volts, the gas mode was set to static, the temperature of ion transfer tube was set to 240 °C). FAIMSPro ion mobility was applied (Standard resolution, total carrier gas flow static at 3.6 liters per minute, and its compensation voltage was set to -45 volts). The isolation window was 20 Th over the mass range.

DIA-LIT 40X10 Turbo auto-injection time

The scan sequence began with an MS1 spectrum (Orbitrap analysis, resolution 120,000, scan range 400–1000 Th, automatic gain control (AGC) target of 300%, maximum injection time 50 ms, RF lens 40%). The precursor's mass range for MS2 analysis (Linear ion trap analysis, *Turbo* scanning mode, auto-injection time) was set from 500 to 900 Th, and the scan range from 200 - 1200 Th. MS2 analysis consisted of higher-energy collisional dissociation (HCD), MS2 AGC was set to 1000%, NCE (normalized collision energy) was 33%, and the nanospray ionization was used in this experiment (positive ion 2300 volts, negative ion 600 volts, the gas mode was set to static, the temperature of ion transfer tube was set to 240 °C). FAIMSPro ion mobility was applied (Standard resolution, total carrier gas flow static at 3.6 liters per minute, and its compensation voltage was set to -45 volts). The isolation window was 10 Th over the mass range.

DIA-LIT 40X10 Turbo 8 ms injection time

The scan sequence began with an MS1 spectrum (Orbitrap analysis, resolution 120,000, scan range 400–1000 Th, automatic gain control (AGC) target of 300%, maximum injection time 50 ms, RF lens 40%). The precursor's mass range for MS2 analysis (Linear ion trap analysis, *Turbo* scanning mode, 8 ms injection time) was set from 500 to 900 Th and the scan range from 200 - 1200 Th. MS2 analysis consisted of higher-energy collisional dissociation (HCD), MS2 AGC was set to 1000%, NCE (normalized collision energy) was 33%, and the nanospray ionization was used in this experiment (positive ion 2300 volts, negative ion 600 volts, the gas mode was set to static, the temperature of ion transfer tube was set to 240 °C). FAIMSPro ion mobility was applied (Standard resolution, total carrier gas flow static at 3.6 liters per minute, and its compensation voltage was set to -45 volts). The isolation window was 10 Th over the mass range.

DIA-LIT 40X10 Rapid auto-injection time

The scan sequence began with an MS1 spectrum (Orbitrap analysis, resolution 120,000, scan range 400–1000 Th, automatic gain control (AGC) target of 300%, maximum injection time 50 ms, RF lens 40%). The precursor's mass range for MS2 analysis (Linear ion trap analysis, *Rapid* scanning mode, auto-injection time) was set from 500 to 900 Th, and the scan range from 200 - 1200 Th. MS2 analysis consisted of higher-energy collisional dissociation (HCD), MS2 AGC was set to 1000%, NCE (normalized collision energy) was 33%, and the nanospray ionization was used in this experiment (positive ion 2300 volts, negative ion 600 volts, the gas mode was set to static, the temperature of ion transfer tube was set to 240 °C). FAIMSPro ion mobility was applied (Standard resolution, total carrier gas flow static at 3.6 liters per minute, and its compensation voltage was set to -45 volts). The isolation window was 10 Th over the mass range.

DIA-LIT 30X13.5 Rapid 30 ms injection time

The scan sequence began with an MS1 spectrum (Orbitrap analysis, resolution 120,000, scan range 400–1000 Th, automatic gain control (AGC) target of 300%, maximum injection time 50 ms, RF lens

40%). The precursor's mass range for MS2 analysis (Linear ion trap analysis, *Rapid* scanning mode, 30 ms injection time) was set from 500 to 900 Th, and the scan range from 200 - 1200 Th. MS2 analysis consisted of higher-energy collisional dissociation (HCD), MS2 AGC was set to 1000%, NCE (normalized collision energy) was 33%, and the nanospray ionization was used in this experiment (positive ion 2300 volts, negative ion 600 volts, the gas mode was set to static, the temperature of ion transfer tube was set to 240 °C). FAIMSPro ion mobility was applied (Standard resolution, total carrier gas flow static at 3.6 liters per minute, and its compensation voltage was set to -45 volts). The isolation window was 13.5 Th over the mass range.

<u>DIA-LIT 25X16 Rapid 31 ms injection time</u>

The scan sequence began with an MS1 spectrum (Orbitrap analysis, resolution 120,000, scan range 400–1000 Th, automatic gain control (AGC) target of 300%, maximum injection time 50 ms, RF lens 40%). The precursor's mass range for MS2 analysis (Linear ion trap analysis, *Rapid* scanning mode, 31 ms injection time) was set from 500 to 900 Th and the scan range from 200 - 1200 Th. MS2 analysis consisted of higher-energy collisional dissociation (HCD), MS2 AGC was set to 1000%, NCE (normalized collision energy) was 33%, and the nanospray ionization was used in this experiment (positive ion 2300 volts, negative ion 600 volts, the gas mode was set to static, the temperature of ion transfer tube was set to 240 °C). FAIMSPro ion mobility was applied (Standard resolution, total carrier gas flow static at 3.6 liters per minute, and its compensation voltage was set to -45 volts). The isolation window was 16 Th over the mass range.

<u>DIA-LIT 34X10 Rapid auto-injection time</u>

The scan sequence began with an MS1 spectrum (Orbitrap analysis, resolution 120,000, scan range 400–1000 Th, automatic gain control (AGC) target of 300%, maximum injection time 50 ms, RF lens 40%). The precursor's mass range for MS2 analysis (Linear ion trap analysis, *Rapid* scanning mode, 31 ms injection time) was set from 500 to 900 Th and the scan range from 200 - 1200 Th. MS2 analysis consisted of higher-energy collisional dissociation (HCD), MS2 AGC was set to 1000%, NCE (normalized collision energy) was 33%, and the nanospray ionization was used in this experiment (positive ion 2300 volts, negative ion 600 volts, the gas mode was set to static, the temperature of ion transfer tube was set to 240 °C). FAIMSPro ion mobility was applied (Standard resolution, total carrier gas flow static at 3.6 liters per minute, and its compensation voltage was set to -45 volts). The isolation window was 11.7 Th over the mass range.

<u>DIA-LIT 45X10 Rapid auto-injection time</u>

The scan sequence began with an MS1 spectrum (Orbitrap analysis, resolution 120,000, scan range 400–1000 Th, automatic gain control (AGC) target of 300%, maximum injection time 50 ms, RF lens 40%). The precursor's mass range for MS2 analysis (Linear ion trap analysis, *Rapid* scanning mode, 31 ms injection time) was set from 500 to 900 Th and the scan range from 200 - 1200 Th. MS2 analysis consisted of higher-energy collisional dissociation (HCD), MS2 AGC was set to 1000%, NCE (normalized collision energy) was 33%, and the nanospray ionization was used in this experiment (positive ion 2300 volts, negative ion 600 volts, the gas mode was set to static, the temperature of ion transfer tube was set to 240 °C). FAIMSPro ion mobility was applied (Standard resolution, total carrier gas flow static at 3.6 liters per minute, and its compensation voltage was set to -45 volts). The isolation window was 9 Th over the mass range.

DIA-LIT 40X10 Normal auto-injection time

The scan sequence began with an MS1 spectrum (Orbitrap analysis, resolution 120,000, scan range 400–1000 Th, automatic gain control (AGC) target of 300%, maximum injection time 50 ms, RF lens 40%). The precursor's mass range for MS2 analysis (Linear ion trap analysis, *Normal* scanning mode, auto-injection time) was set from 500 to 900 Th and the scan range from 200 - 1200 Th. MS2 analysis consisted of higher-energy collisional dissociation (HCD), MS2 AGC was set to 1000%, NCE (normalized collision energy) was 33%, and the nanospray ionization was used in this experiment (positive ion 2300 volts, negative ion 600 volts, the gas mode was set to static, the temperature of ion transfer tube was set to 240 °C). FAIMSPro ion mobility was applied (Standard resolution, total carrier gas flow static at 3.6 liters per minute, and its compensation voltage was set to -45 volts). The isolation window was 10 Th over the mass range.

DIA-LIT 30X10 Normal 50 ms injection time

The scan sequence began with an MS1 spectrum (Orbitrap analysis, resolution 120,000, scan range 400–1000 Th, automatic gain control (AGC) target of 300%, maximum injection time 50 ms, RF lens 40%). The precursor's mass range for MS2 analysis (Linear ion trap analysis, *Normal* scanning mode, auto-injection time) was set from 500 to 900 Th and the scan range from 200 - 1200 Th. MS2 analysis consisted of higher-energy collisional dissociation (HCD), MS2 AGC was set to 1000%, NCE (normalized collision energy) was 33%, and the nanospray ionization was used in this experiment (positive ion 2300 volts, negative ion 600 volts, the gas mode was set to static, the temperature of ion transfer tube was set to 240 °C). FAIMSPro ion mobility was applied (Standard resolution, total carrier gas flow static at 3.6 liters per minute, and its compensation voltage was set to -45 volts). The isolation window was 13.5 Th over the mass range.

DIA-LIT 20X10 Normal 76 ms injection time

The scan sequence began with an MS1 spectrum (Orbitrap analysis, resolution 120,000, scan range 400–1000 Th, automatic gain control (AGC) target of 300%, maximum injection time 50 ms, RF lens 40%). The precursor's mass range for MS2 analysis (Linear ion trap analysis, *Normal* scanning mode, auto-injection time) was set from 500 to 900 Th and the scan range from 200 - 1200 Th. MS2 analysis consisted of higher-energy collisional dissociation (HCD), MS2 AGC was set to 1000%, NCE (normalized collision energy) was 33%, and the nanospray ionization was used in this experiment (positive ion 2300 volts, negative ion 600 volts, the gas mode was set to static, the temperature of ion transfer tube was set to 240 °C). FAIMSPro ion mobility was applied (Standard resolution, total carrier gas flow static at 3.6 liters per minute, and its compensation voltage was set to -45 volts). The isolation window was 20 Th over the mass range.

DIA-LIT 40X10 Enhanced auto-injection time

The scan sequence began with an MS1 spectrum (Orbitrap analysis, resolution 120,000, scan range 400–1000 Th, automatic gain control (AGC) target of 300%, maximum injection time 50 ms, RF lens 40%). The precursor's mass range for MS2 analysis (Linear ion trap analysis, *Enhanced* scanning mode, auto-injection time) was set from 500 to 900 Th, and the scan range from 200 - 1200 Th. MS2 analysis consisted of higher-energy collisional dissociation (HCD), MS2 AGC was set to 1000%, NCE (normalized collision energy) was 33%, and the nanospray ionization was used in this experiment (positive ion 2300 volts, negative ion 600 volts, the gas mode was set to static, the temperature of ion transfer tube was set to 240 °C). FAIMSPro ion mobility was applied (Standard resolution, total carrier gas flow static at 3.6 liters per minute, and its compensation voltage was set to -45 volts). The isolation window was 10 Th over the mass range.
